# Selective peak inference: Unbiased estimation of raw and standardized effect size at local maxima

**DOI:** 10.1101/500512

**Authors:** Samuel J. Davenport, Thomas E. Nichols

## Abstract

The spatial signals in neuroimaging mass univariate analyses can be characterized in a number of ways, but one widely used approach is peak inference: the identification of peaks in the image. Peak locations and magnitudes provide a useful summary of activation and are routinely reported, however, the magnitudes reflect selection bias as these points have both survived a threshold and are local maxima. In this paper we propose the use of resampling methods to estimate and correct this bias in order to estimate both the raw units change as well as standardized effect size measured with Cohen’s *d* and partial *R*^2^. We evaluate our method with a massive open dataset, and discuss how the corrected estimates can be used to perform power analyses.

## 1 Introduction

Any time a set of noisy data is scanned for the largest value, this value will be an overestimate of the true, noise-free maximum. This effect is known as regression to the mean or the winner’s curse and occurs because, at random, some of the variables get lucky and take on high values. In neuroimaging, an analysis produces a test statistic at each voxel, and there are then a number of inference methods available to assess the evidence for a true activation. Voxel, peak and cluster level inference are the most common.^1^ When papers report the effect size at a peak it is biased due to the winner’s curse. This bias is typically caused by two factors, firstly the observed peaks have been chosen such that they lie above a threshold and secondly the value at each peak is the largest value in a local region around the peak (though any type of threshold or selection of the peaks based on their magnitude or a another correlated quantity has the potential to cause them to be biased). In order to determine the true effect sizes we have to account for this bias.

This issue is well-known in neuroimaging and is called circular inference or double dipping (Kriegeskorte et al., 2009). Vul et al. (2009) conducted a review and found the problem to be widespread in the fMRI literature, to much controversy. In their metaanalysis of 55 articles, where the test-statistic at each voxel was the correlation between %BOLD signal and a personality measure, they found that correlations observed were spuriously high in papers that reported values at peaks, reflecting a bias due to the winner’s curse.

The main existing solution to this problem in neuroimaging is data-splitting, where the first half of the data is used to find significant regions and the other half is used to calculate effect sizes; Kriegeskorte et al. (2010), Kriegeskorte et al. (2009). While this produces unbiased values, the estimates have larger variance as they are calculated using only half of the data. For the same reason, the locations of local maxima will be less accurate than if they had been calculated using the whole dataset. These problems are especially serious when the sample sizes are small. A widely used alternative is to select a voxel or ROI a priori based on previous studies and to only calculate the effect at that location. While this approach is unbiased it has the disadvantage that only the pre-specified voxel or ROI can be considered, and not the peaks found in the observed data. Instead if it were possible to use all of the data to estimate locations and effect sizes whilst still obtaining unbiased point estimates of the signal magnitude we would obtain much more accurate estimates of the peak locations. This type of approach, where you use all of the data, is known as post-model selection or selective inference and has recently generated a lot of interest; see Berk et al. (2013), Lee et al. (2016) and in particular Taylor and Tibshirani (2015) for a good overview.

A similar problem arises in genetics, see Göring et al. (2001), and there has been much recent work on correcting for selection in this setting. Zhong and Prentice (2008), Ghosh et al. (2008) and Xiao and Boehnke (2011) consider pointwise correction by calculating the distribution of the effect statistic conditional on it being significant while Zhou and Wright (2015), Sun and Bull (2005), Wu et al. (2006), Yu et al. (2007) and Jeffries (2006) consider resampling based approaches. In the imaging literature, Rosenblatt and Benjamini (2014) propose a selective inference approach to obtain unbiased confidence intervals but not point estimates. Under the assumption of constant variance Benjamini and Meir (2014) propose a method to correct all voxels above a threshold, analogous to the genetics pointwise correction discussed above. However, this doesn’t take account of the effect of selecting peaks or the dependence between voxels. Esterman et al. (2010) use a leave one out cross validation approach to provide corrected estimates however this approach has the disadvantage that each instance of resampled data has a different estimate of the significant locations meaning that these are not identifiable. We employ a bootstrap resampling method that provides point estimates of local maxima, accounting for both the peak height and the location within the image. We use all of the data to determine significant locations, meaning that these locations are consistent across resamples and relate to the original statistic image used for inference.

The idea of using an estimate other than the sample mean to provide an estimate for the mean is first due to Stein (1956) and James and Stein (1961) who introduced the famous James-Stein estimator. More recently there has been work to correct for the bias in estimating the means of the largest observed values of a given distribution. Efron (2011) uses an empirical Bayes technique to correct for this bias, an approach that has been applied in the genetics literature (Ferguson et al. (2013)). In the case of independent random variables that each come from distributions belonging to a known parametric family, Simon and Simon (2013) introduced a frequentist method to correct bias and Reid et al. (2014) details a post-model selection approach which involves calculating the distribution of Gaussian random variables conditional on being selected.

Brain imaging data is more complicated than these other settings as it has complex spatial and temporal dependencies. However, for group analyses we can take advantage of the fact that data from different subjects is independent. This allows us to employ a bootstrap approach to resample the data while preserving the spatial dependence structure. Our approach is based on an extension of Simon and Simon (2013) to account for dependence proposed by Tan et al. (2014), where a nonparametric bootstrap is used to estimate bias in effect sizes, motivated by a genetics application. We provide a detailed framework for this method and show how it can be applied in the context of neuroimaging. The novel contribution of our work is to develop point estimates which account for selective inference bias due to thresholding and the use of local maxima. We develop these methods to obtain accurate estimates of Cohen’s *d* and *R*^2^, two quantities that are essential for power analyses. See Mumford (2012) for an overview and Appendix D for the mathematical details of how to use power analyses in neuroimaging.

We use functional and structural magnetic resonance images (MRI) from 8,940 subjects from the UK biobank. The size of this dataset allows us to validate our methods in a way that has never been possible before the availability of data of such scale, allowing us to set aside 4,000 subjects to provide an accurate estimate of the truth and divide the remaining subjects into small groups in order to test our methods. The importance of these sort of real data empirical validations is highlighted by recent work on the validity of cluster size inference (Eklund et al., 2016).

The structure of this paper is as follows. Section 2 explains the details behind the bootstrapping method and how it can be applied to one-sample and the more general linear model scenario. In the one-sample case our method provides corrected estimates of the raw effect (e.g. %BOLD mean where BOLD stands for the Blood-oxygen-level-dependent signal) and Cohen’s *d* at the locations of peaks of the one-sample *t*-statistic found to be significant after correction for multiple comparisons. In the case of the general linear model it provides corrected estimates of partial *R*^2^ values. Section 2.4 discusses the methods used for big data evaluation. Section 3.1 illustrates the methods on simulated data and Section 3.2 applies the techniques to one-sample analysis of functional imaging data and GLM analysis of structural gray matter data obtained using voxel based morphometry (VBM). In Section 3.3 we apply our method to a dataset from the Human Connectome Project that involves a contrast for working memory and obtain corrected Cohen’s *d* and %BOLD values at significant peaks.

## 2 Methods

Let 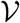 be the set of voxel locations corresponding to the brain or some subset under study and define an **image** to be a map which takes voxel locations to intensities. Given an image *Z* and a connectivity criterion that determines the neighbours of each voxel (we use a connectivity criterion of 18 in our 3D analyses), define a **local maxima or peak** of *Z* to be a voxel such that the value that *Z* takes at that location is larger than the value *Z* takes at neighbouring voxels; see Appendix C for a rigorous definition of this.

### 2.1 One-Sample

Suppose that we have *N* subjects and for each *n* = 1, …, *N* a corresponding random image *Y_n_* on 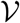 such that for every voxel 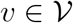,

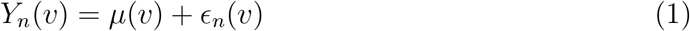

where *μ*(*v*) is the common mean intensity, and the noise terms *ϵ*_1_, …, *ϵ_n_* are iid mean zero random images from some unknown multivariate distribution on 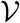. Let 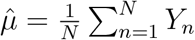 be the sample mean image and let 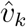 be the location of the *k*th largest local maximum of 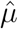 above a screening threshold *u*. For each *k*, we are interested in inferring on 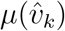, the value of *μ* at the location 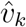, improving on the biased circular estimate 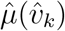.

#### 2.1.1 Peak Estimation

The noise distribution in model (1) is unknown so in order to estimate the bias of 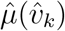 we use the data to generate bootstrap samples without making any distributional assumptions. This allows us to obtain an estimate of the bias for each bootstrap iteration as in Tan et al. (2014). For each maxima 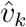 we estimate the bias-corrected value as 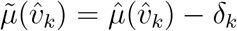, where *δ_k_* are bias correction terms found as described in Algorithm 1 below. See Table 1 for a variable key.

**Table 1:** Variable Key: Note that for clarity we have dropped the index *v* above, all operations are done pointwise on the images at each of their voxels.

| Variable | Definition |
| --- | --- |
| $\hat{\mu}, \hat{\mu}_b$ | $\hat{\mu} = \frac{1}{N} \sum_{i=1}^N Y_n$ and $\hat{\mu}_b$ is the $b$th bootstrapped version of $\hat{\mu}$. |
| $\hat{\sigma}^2, \hat{\sigma}_b^2$ | $\hat{\sigma}^2 = \frac{1}{N-1} \sum_{n=1}^N (Y_n - \hat{\mu})^2$ and $\hat{\sigma}_b^2$ is the $b$th bootstrapped version of $\hat{\mu}$. |
| $\hat{d}, \hat{d}_b$ | $\hat{d} = \hat{\mu}/\hat{\sigma}$ and $\hat{d}_b = \hat{\mu}_b/\hat{\sigma}_b$ |
| $R^2, R_b^2$ | $R^2$ is the partial $R^2$ image and $R_b^2$ is the $b$th bootstrapped version of $R^2$. |
| $\hat{\delta}_{k,b}$ | $\hat{\delta}_{k,b}$ is the $b$th bootstrap estimate of the bias at the $k$th largest peak. |
| $\hat{\delta}_k$ | $\hat{\delta}_k = \frac{1}{B} \sum_{b=1}^B \hat{\delta}_{k,b}$ is the estimate of the bias at the $k$th largest peak. |
| $\hat{v}_k$ | $\hat{v}_k$ is the location of the $k$th largest peak in the observed effectsize image i.e. $\hat{\mu}, \hat{d}, R^2$ respectively. |
| $\hat{v}_{k,b}$ | $\hat{v}_{k,b}$ is the location of the $k$th largest peak in the $b$th bootstrap image i.e. $\hat{\mu}_b, \hat{d}_b, R_b^2$ respectively. |

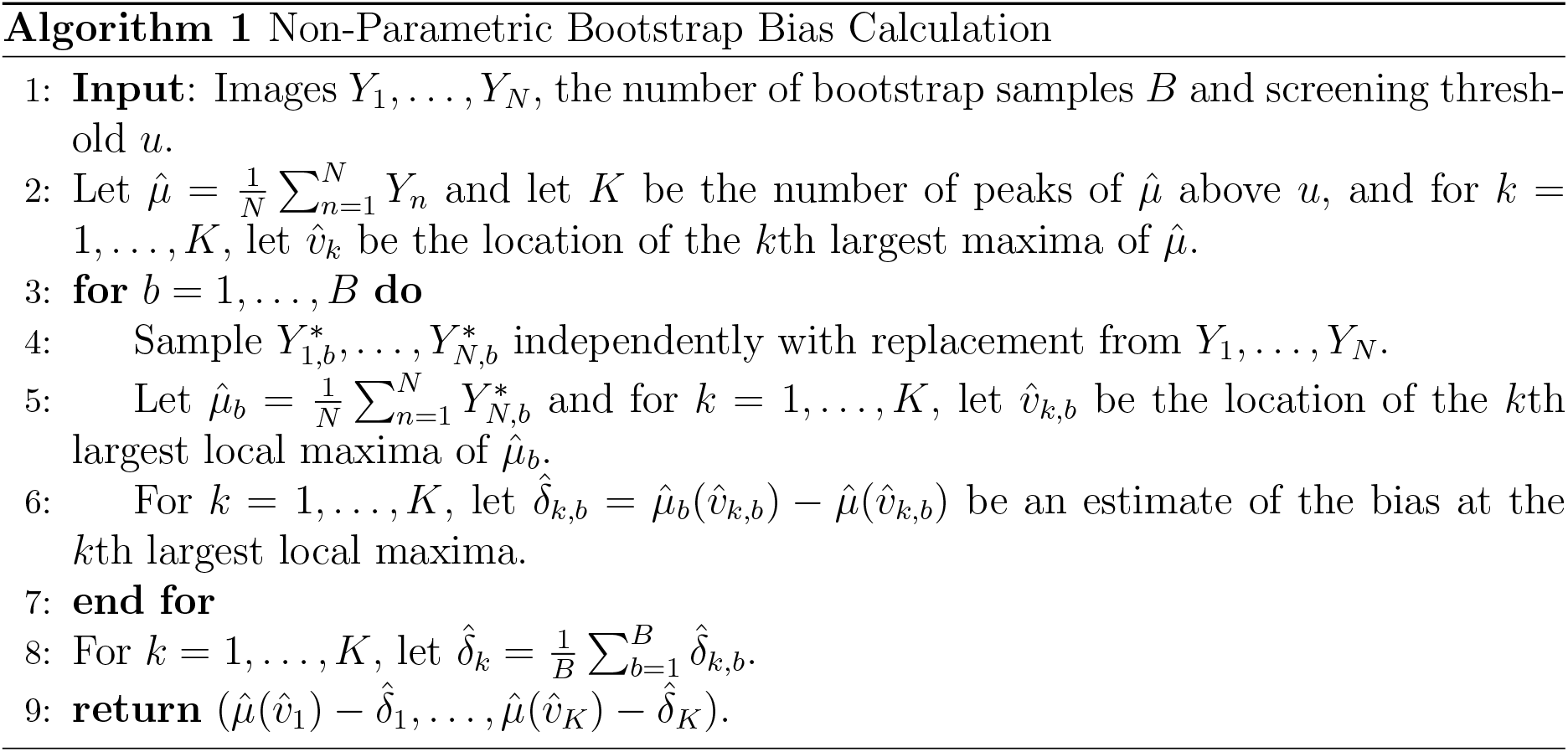

Figure 1 provides an illustrative 1D simulation on a grid of 160 voxels where we consider the case *k* = 1: the global maximum. Here, *N* = 20 and for each *n* = 1, …, *N* the error images are created by simulating iid Gaussians at each voxel with variance 4 and then smoothing this with a 6 voxel FWHM Gaussian kernel. The bias above the noise-free signal (*δ*_1_) is evident, and is estimated by comparing a bootstrap sample to the original, yielding an estimate of 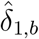. *δ*_1_ is the bias of the empirical mean relative to the true mean.

**Figure 1:**
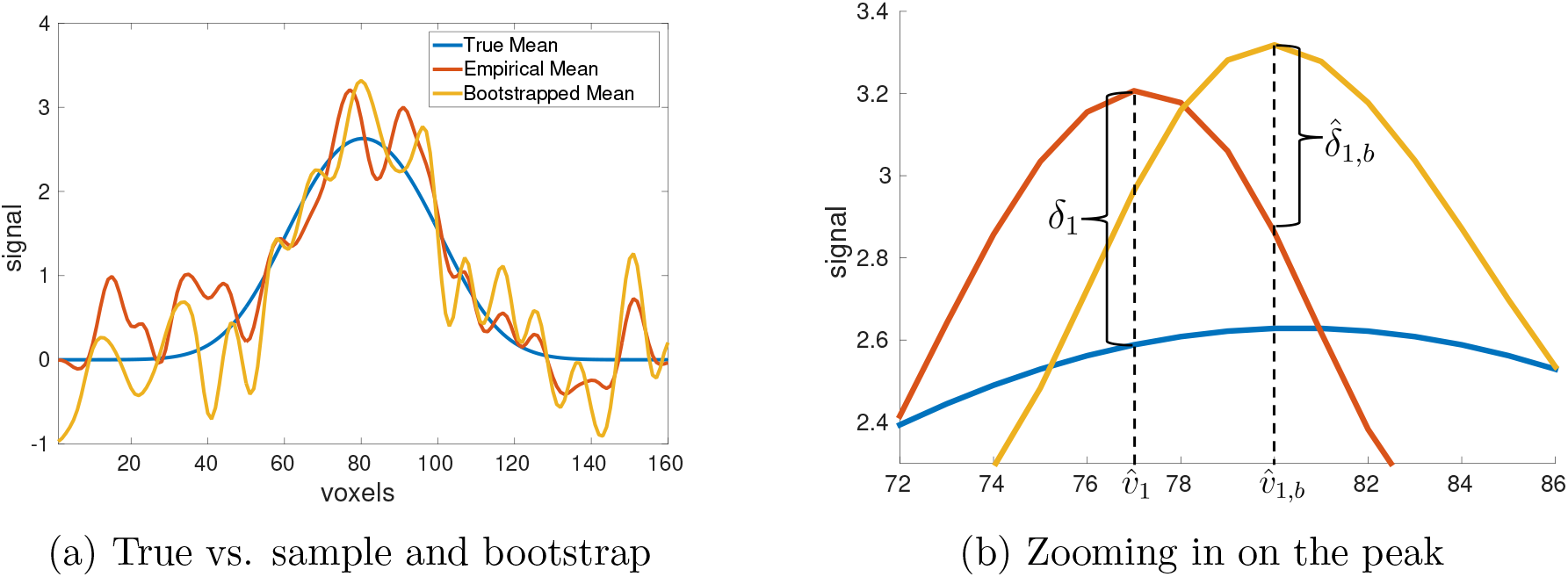
Illustration of our bootstrap peak bias correction method on a simple annotated example. Here our set of voxels is 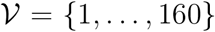, the true mean *μ* is shown in blue, the empirical mean 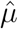 is shown in red and one sample bootstrap realization (iteration *b*) is shown in yellow. The figure on the right is a zoomed in version of the figure on the left from voxel 72 to 86. The top peak is biased above the true value by *δ*_1_. Using this realization the height of the bootstrapped peak is compared to the height of the empirical mean at the location of the peak 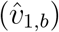, resulting in an estimate 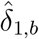 of the bias.

#### 2.1.2 Peak Estimation for Effect Size

While the above method is based on the sample mean, neuroimaging studies typically base their inferences on statistic images. In the simplest setting, a one-sample analysis of fMRI contrast data, we are testing

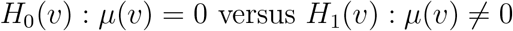

at each 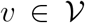, using the statistic image, in order to determine whether there is an activation at that voxel. Given the voxels that have been determined to be active we are interested in estimating two different quantities, the effect size (measured in terms of Cohen’s *d*) and the raw unit, i.e. %BOLD change for fMRI.

We first need to define the test-statistic image. Define *σ*^2^ to be the population variance image, estimated in an unbiased manner using

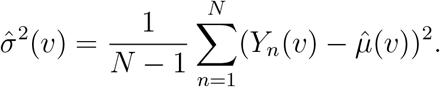

In order to perform one-sample hypothesis testing, the *t*-statistic

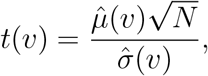

is typically used. If the data are Gaussian at each voxel this follows a *t*-distribution with *N* – 1 degrees of freedom.

For a voxel *v*, *H*_0_(*v*) is rejected if *t*(*v*) lies above a screening threshold *u*. While a threshold *u* on a mean image is ultimately arbitrary, on a statistic image we can choose a value of *u* to control false positives at a desired level while controlling for multiple testing. For example, we can use results from the theory of random fields to find a *u* such that the familywise error rate, the chance of one or more false positives over the image, is controlled; Worsley et al. (1996), Friston et al. (1994).

While ubiquitously reported, *t*-statistic values are not interpretable across studies, as they depend on the sample size and grow to infinity with *N*. Good practice, and in particular to facilitate power analyses (see Appendix D), requires computation of a standardized effect size such as Cohen’s *d*, which at each voxel *v* is defined as

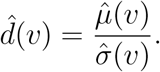

As this is just the one-sample *t*-statistic divided by 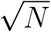, the peaks in the *t*-statistic image will be at the same locations as those of the one-sample Cohen’s *d*. Algorithm 2 describes how to compute bias-corrected estimates of Cohen’s *d* peaks, which we evaluate with simulated data (Section 3.1.1) and real task fMRI data (Section 3.2.1). The one-sample Cohen’s *d* is a biased estimator for the population Cohen’s *d*:

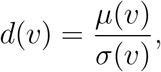

with 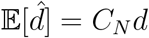 where *C_N_* is a correction factor that depends on the degrees of freedom. *C*_N_ → 1 as *n* → ∞ but for finite samples we need to account for it; see Appendix D for details.

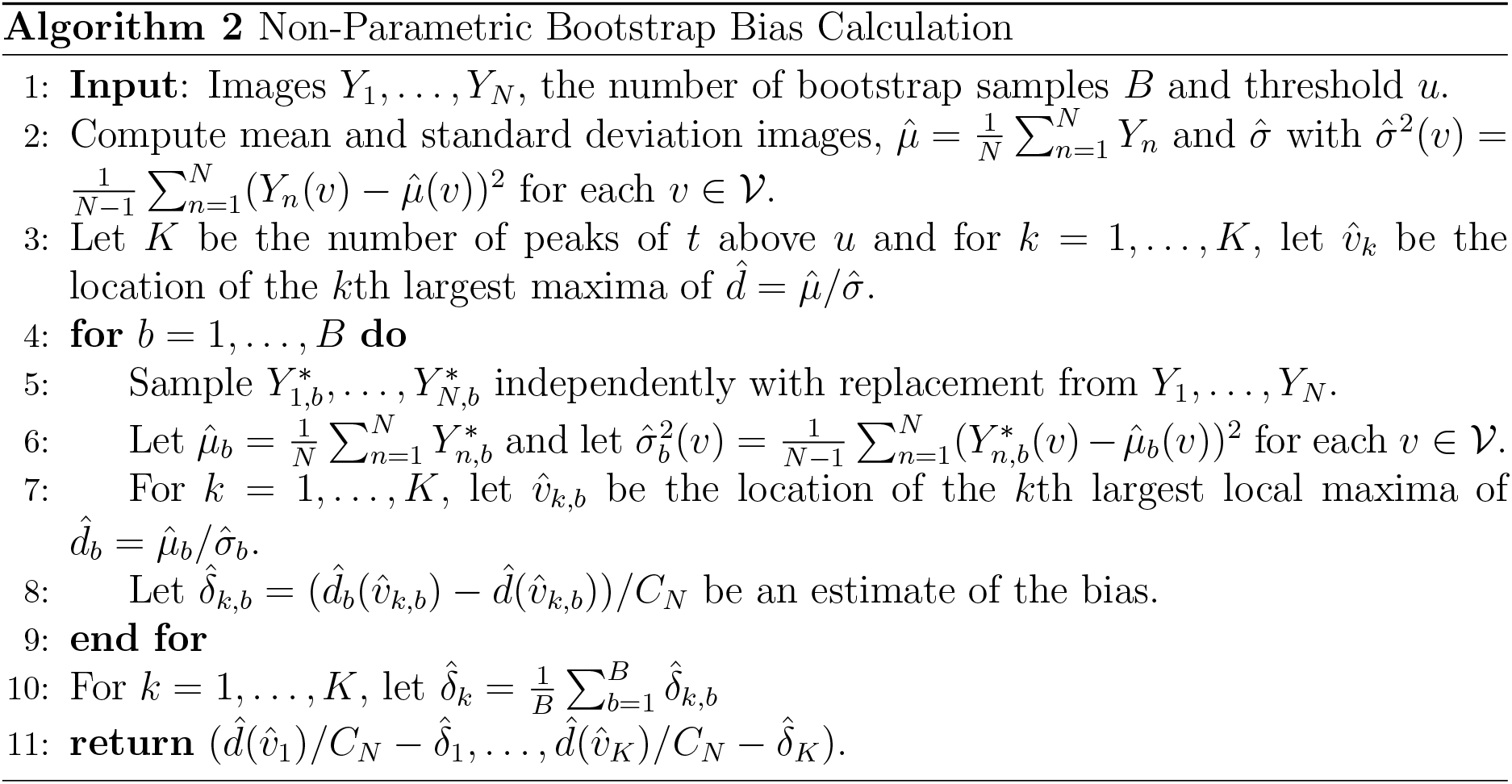

#### 2.1.3 Estimation of the Mean at the Location of Effect Size Peaks

In fMRI the underlying mean *μ* corresponds to the true %BOLD signal which is the expected value of the contrast image for each subject, see Mumford and Nichols (2009) for more details. We asssume that first level models have been fit to give contrast images for each subject. At the first level a number of pre-processing steps are implemented including registration, motion correction, and normalization which affect the %BOLD signal. Some authors, Chen et al. (2017) most recently, have argued that the attention given to statistic images is misguided, and more focus should be given to results with interpretable units, i.e. %BOLD. In order to estimate the mean while still controlling for false positives one needs to use the *t*-statistic image to identify significant peaks and then estimate the raw unit (e.g. %BOLD) change at these locations. This is easily accomplished with a small modification to Algorithm 2, computing in Step 8 instead a bias in raw effect units of:

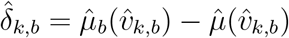

and returning 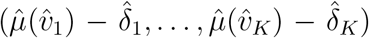 instead. See Section 3.1.2 for simulated evaluations of this approach and Section 3.2.2 for validation of this approach on the estimation of %BOLD mean at local maxima of the *t*-statistics of task fMRI data.

#### 2.1.4 Existing One-Sample Methods

We compare the bootstrap approach to circular inference (no correction) and data-splitting, the main two approaches used in the literature. After finding the number of peaks above the threshold as in Algorithm 2, the circular inference uncorrected estimates are simply 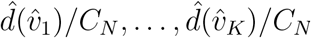.

Data-splitting proceeds as follows. First we divide the images into two groups: *Y*_1_, …, *Y*_*N*/2_ and *Y*_*N*/2+1_, …, *Y_N_*. Let 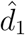 and 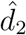 be the image estimates of Cohen’s *d* from the first and second half of the subjects respectively. Using a threshold *u* we find the peaks of the one-sample *t*-statistic 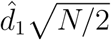 that lie above *u*, at locations 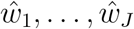 for some number of peaks *J* (note that *u* must be adjusted to account for the fact we are using half the data). The data-splitting estimates of the peak values are 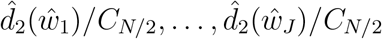. See Figure 4 for an illustration of the different methods applied to a sample consisting of 50 subjects. Note that in general the number of significant peaks found by data-splitting will be lower than the number found using all of the data as with half the number of subjects there is less power to detect activation.

### 2.2 General Linear Model

Having introduced the method in the simplified setting of a one-sample model, we now turn to the regression setting. Here, we will often have no practical meaningful units; for example, for a covariate of age, the units of the coefficient are clear (expected change in response per year) but awkward, and more typically users will want to reference the partial coefficient of determination, partial *R*^2^: the proportion of variance explained by one (or more) predictors not already explained by other terms in the model. Hence we now generalize our method to obtain corrected estimates of the peak partial *R*^2^.

Let *Y* be an *N*-dimensional random image such that for each 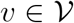, we assume the following linear model,

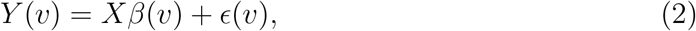

for an *N* × *p* design matrix *X* and a parameter vector 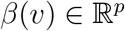 where *ϵ* is the random image of the noise such that *ϵ*(*v*) = (*ϵ*_1_(*v*), …, *ϵ_N_*(*v*))^*T*^ for each 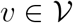 (where the *ϵ_i_* are iid zero mean zero random images). Then we are interested in testing

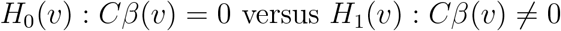

for some contrast matrix 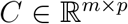 where *m* is the number of contrasts that we simultaneously test. We can test this at each voxel with the usual *F*-test, which at each voxel *v* is

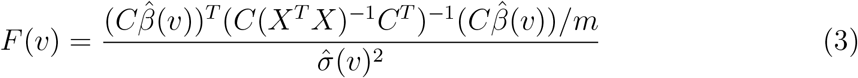

where 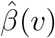 is the least squares estimate of *β*(*v*) and 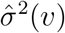 is the estimate of the error variance at each voxel. Then assuming normality of *ϵ*, under the null hypothesis *H*_0_(*v*), *F*(*v*) has an *F_m,N–p_* distribution and can therefore be used for testing purposes. We will incorporate this into our bootstrap algorithm in order to establish which peaks are significant.

Define *R*^2^ be the image with the estimated partial *R*^2^ values for comparing the null model against the alternative at each voxel; we then seek a bias corrected estimate of the partial *R*^2^ at local maxima. See Appendix A for details on how partial *R*^2^ is formally defined. Bootstrapping in the general linear model scenario is based on the residuals; see Davison et al. (2003) Chapter 6. This leads to Algorithm 3.

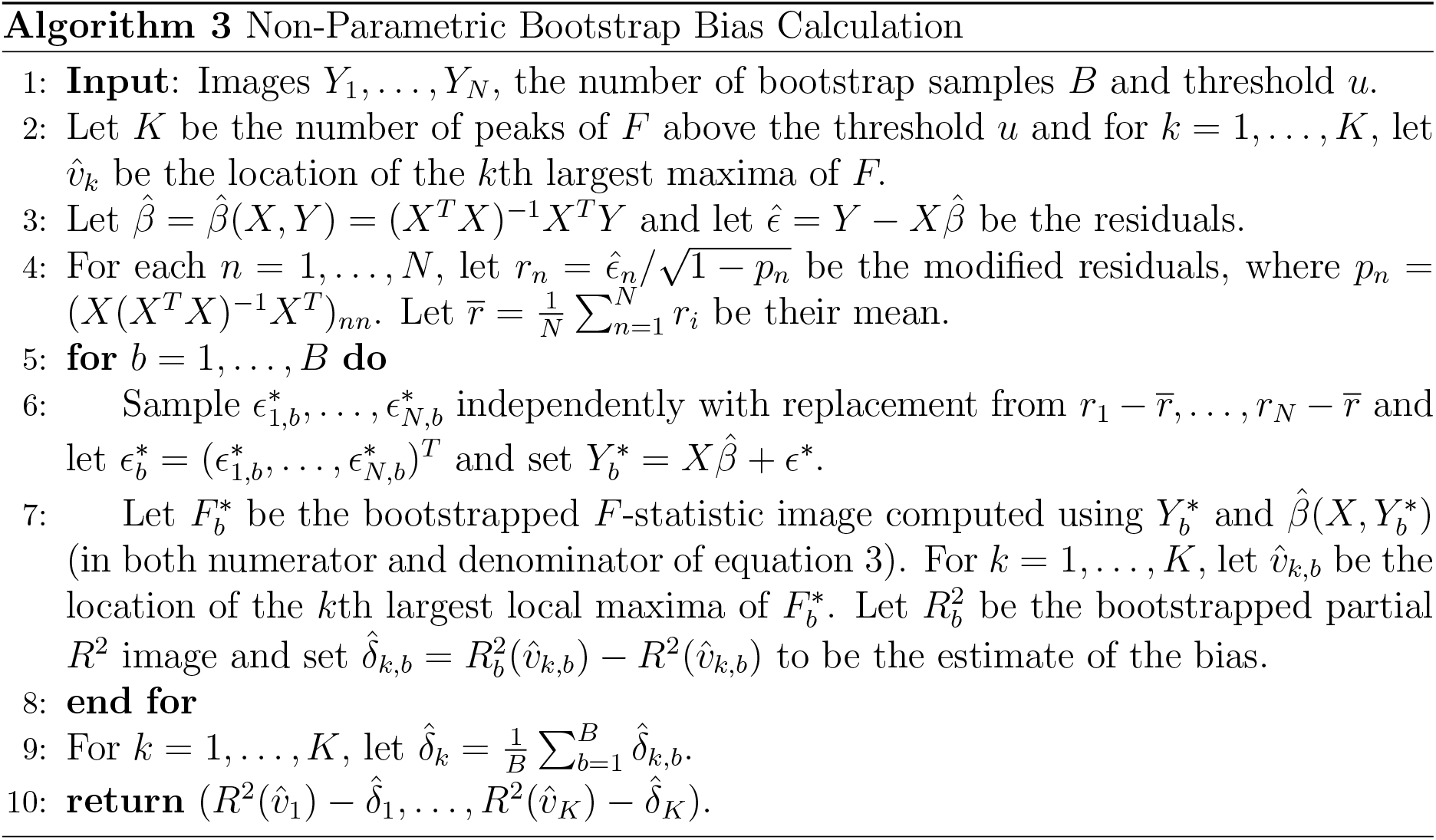

In fMRI we are often interested in the case where 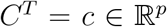 is a contrast vector in which case we can also test using the *t*–statistic

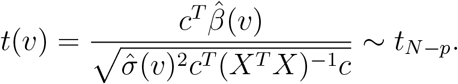

which allows us to perform either one or two sided tests in order to determine significance before bootstrapping.

As in the Section 2.1.4 we can define circular inference and data-splitting estimates. See Section 3.2.3 for validation of the use of the bootstrap and comparisons between the methods in a GLM scenario where gray matter images are regressed against the age of the participants and an intercept. Note that there is no analogous correction factor *C_N_* for *R*^2^ and so even the data-splitting estimates will not be completely unbiased as estimates for the population *R*^2^. However as can be seen from implementation of the algorithms in simulations (see Supplementary Material Figures S2, S3) and on real data this bias is comparatively small.

### 2.3 Simulations

#### 2.3.1 One Sample Mean

In order to test Algorithm 1 we generate 3D simulations on a 91 × 109 × 91 size grid which makes up our set of voxels 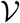. This grid size is that which results from using MNI space and 2mm voxels. We generate data according to model (1) with underlying mean consisting of 3 different peaks with magnitudes of 2, 4, 4 placed at different points of the image. For the *ϵ_n_* we use mean zero Gaussian noise smoothed with an FWHM of 3 voxels, scaled to have variance 1. In order to evaluate the methods we consider sample sizes of *N* = 20, 30, …, 100 and for each sample size generate 1000 realizations. We use a threshold *u* = 2, this value has been chosen arbitrarily, in practice it could be chosen based on domain knowledge about the underlying signal.

#### 2.3.2 One Sample

In order to test the performance of Algorithm 2 for estimation of Cohen’s *d* and the mean we generate 3D simulations (on the same set of voxels 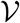 as above) according to model (1) with underlying mean consisting of 9 different peaks each with magnitude 1/2, with one located near corner and one at the centre of the image. See Figure 2 for a slice through this signal and one realization. For the *ϵ_n_* we use mean zero Gaussian noise smoothed with an FWHM of 6mm, scaled to have variance 1.

**Figure 2:**
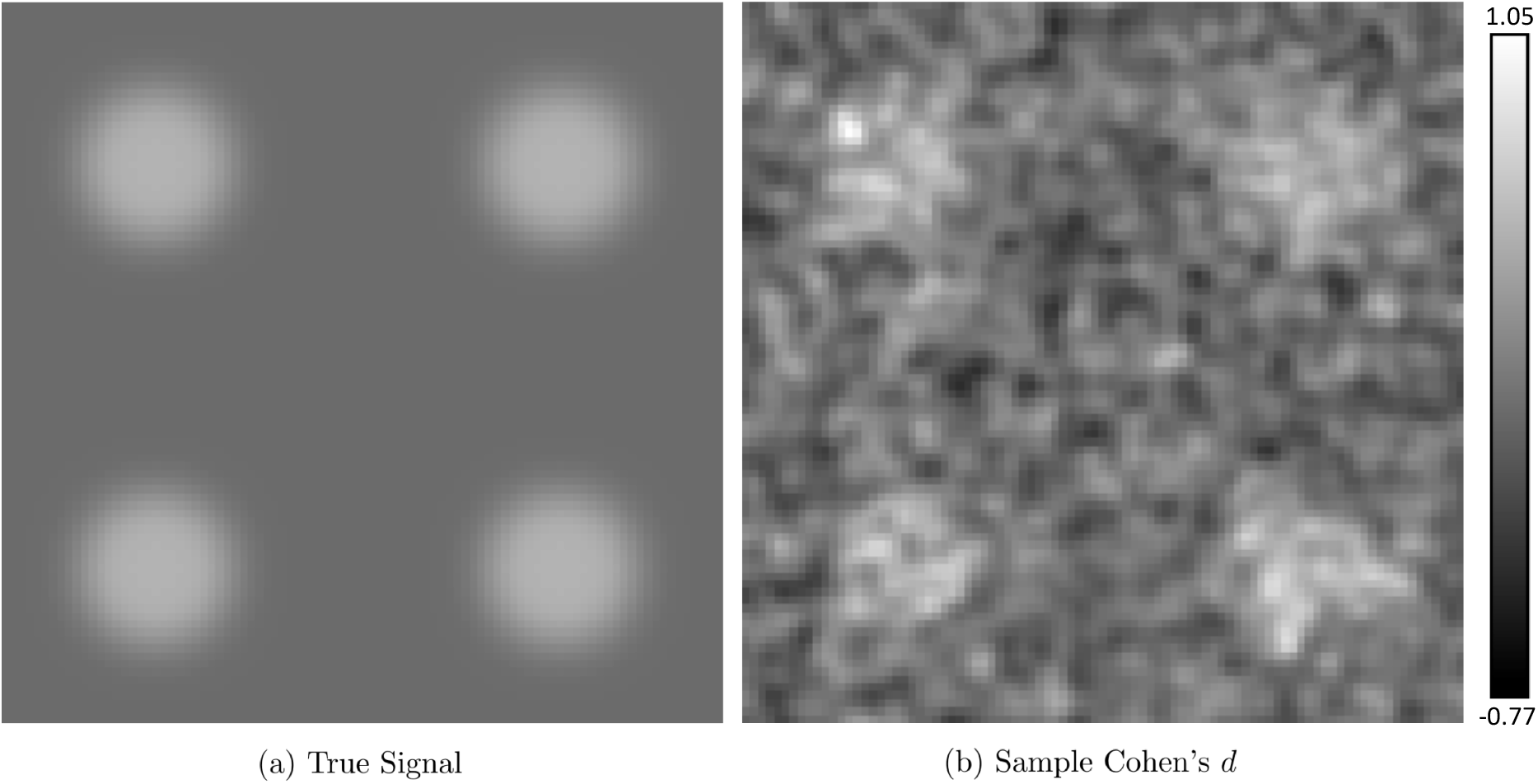
Simulation true signal and one realization. Panel (a) illustrates a slice through the true signal corresponding to the plane *y* = 20, with maximum intensity *d* = 1/2. Panel (b) illustrates the same slice through the one sample Cohen’s *d* for 50 subjects. Data for each subject is computed by adding Gaussian noise (with 3 voxel FWHM) to the signal.

In order to evaluate how the methods compare as the sample size increases we consider *N* random images, *N* = {20, 30, …, 100}, generating 1,000 realizations (of the simulations described above) for each *N*. We use an additional simulation to find the voxelwise threshold that controls the familywise error rate at 5%; for each *N* we generate 5,000 null *t*_*N*–1_ random fields (computed by taking the one-sample *t*-statistic of *N* zero mean Gaussian random fields with 3 voxel FWHM) and take the 95% quantile of the distribution of the maximum.

In order to evaluate how the methods compare as the variance changes we generate 1000 realizations (for each realization we generate 50 subjects) and adjust the variance (which is constant over the image) such that the peak Cohen’s *d* takes the values: {0.1, 0.2, …, 0.7} rather than just 0.5. We controlled the voxelwise familywise error rate as described above.

#### 2.3.3 General Linear Model

In order to test the performance of Algorithm 3 for estimation of partial *R*^2^ we generate 3D simulations (on the same set of voxels 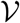 as above) according to the model

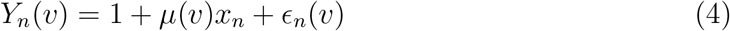

where 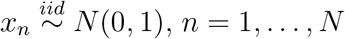 and the *ϵ_n_* are iid random images which are mean zero Gaussian, with 3 voxel FWHM and scaled to have variance 1. *μ* consists of 9 different peaks each with magnitude 0.5822, with one located near corner and one at the centre of the image. The value 0.5822 has been chosen so that the power matches that of the one-sample simulations, see Supplementary Material Section S2 for details. As for the one-sample simulations, for *N* ∈ {20, …, 100} we generate 1,000 realizations of the above model and calculate a voxelwise threshold using additional simulations.

In order to evaluate how the methods compare as the variance changes we generate 1000 realizations and change the variance (which is constant over the image) such that the peak *R*^2^ takes the values: {0.1, 0.2, …, 0.6}. We consider *N* = 50 and *N* = 100 in order to illustrate what happens when you have a sufficiently large number of subjects. We controlled the voxelwise familywise error rate as described above.

### 2.4 Big Data Validation

In order to test our methods we take advantage of the large sample sizes in the UK Biobank. This enables us to set aside 4,000 (randomly selected) subjects in order to compute a very accurate estimate of the mean, Cohen’s *d* or partial *R*^2^ value. We will refer to this 4,000-subject estimate of the effect as the ground truth. See Appendix B for details on how the ground truth is computed in the different settings. Implementing large linear models with missing data can be computationally burdensome so we outline efficient methods for dealing with this in Appendices B.3 and B.4.

We divide the remaining 4,940 subjects into groups similar in size to those used in typical fMRI/VBM studies. For each such group we apply all three methods and compare the values obtained to the ground truth calculated using the 4000 subjects, allowing the performance of the methods across groups to be evaluated. In each small sample, we consider only complete-data voxels, as is typical in neuroimaging analyses.

#### 2.4.1 Image Acquisition

The UK Biobank is a prospective epidemiological resource combining questionnaires, physical and cognitive measures, and biological samples in a sample of 500,000 subjects in the United Kingdom, aged 40-69 years of age at baseline recruitment. The UK Biobank Imaging Extension provides extensive MRI data of the brain, ultimately on 100,000 subjects. We use the prepared data available from the UK Biobank; full details on imaging acquisition and processing can be found in Miller et al. (2016), Alfaro-Almagro et al. (2018) and from UK Biobank Showcase^2^; a brief description is provided here.

The task fMRI data uses the block-design Hariri faces/shapes task Hariri et al. (2002), where the participants are shown triplets of fearful expressions and, in the control condition, triplets of shapes, and for each event perform a matching task. A total of 332 T2*-weighted blood-oxygen level-dependent (BOLD) echo planar images were acquired in each run [TR=0.735s, TE=39ms, FA=52°, 2.4mm^3^ isotropic voxels in 88 × 88 × 64 matrix, ×8 multislice acceleration]. Standard preprocessing and task fMRI modeling was conducted in FEAT (FMRI Expert Analysis Tool); part of the FSL software http://www.fmrib.ox.ac.uk/fsl). After head-motion correction and Gaussian kernel of FWHM 5mm, a linear model was fit at each voxel resulting in contrast images for each subject.

Structural T1-weighted images were acquired on each subject [3D MPRAGE, 1mm^3^ isotopic voxels in 208 × 256 × 256 matrix]. Images were defaced and nonlinearly warped to MNI152 space using FNIRT (FMRIB’s Nonlinear Image Registration Tool). For VBM, tissue segmentation was performed with FSL’s FAST (FMRIB’s Automated Segmentation Tool), producing images of gray matter that were subsequently warped to MNI152 space, and modulated by the Jacobian of the warp field. Warped modulated images were written with voxel sizes of 2mm.

Additional processing consisted of transforming intrasubject contrast maps to MNI space with 2mm using nonlinear warping determined by the T1 image and an affine registration of the T2* to the T1 image. We additionally apply a smoothing of 3mm FWHM to the modulated gray matter images.

#### 2.4.2 Task fMRI analysis

We have faces-shapes contrast images from 8,940 subjects and consider the mean and one sample Cohen’s *d*. We compute the 4,000-subject Cohen’s *d* ground truth image for voxels with data for at least 100 subjects (Figure 3). For a given sample size *N*, let *G_N_* = [4940/*N*] be the number of groups of size *N* into which we can divide the 4,940 remaining subjects^3^. This division enables a comparison of the performance of the three available methods, circular inference, data-splitting and the bootstrap. As in the simulations, we measure the performance in terms of bias, standard deviation and root mean squared error (RMSE) as defined in Section 2.5. We use sample sizes of 20, 50 and 100 to illustrate the performance of the methods, these sizes have been chosen since they are representative of those typically used in fMRI. Sections 3.2.1 and 3.2.2 present the results of applying the methods and Figure 4 illustrates these methods applied to an exemplar sample consisting of 50 subjects.

**Figure 3:**
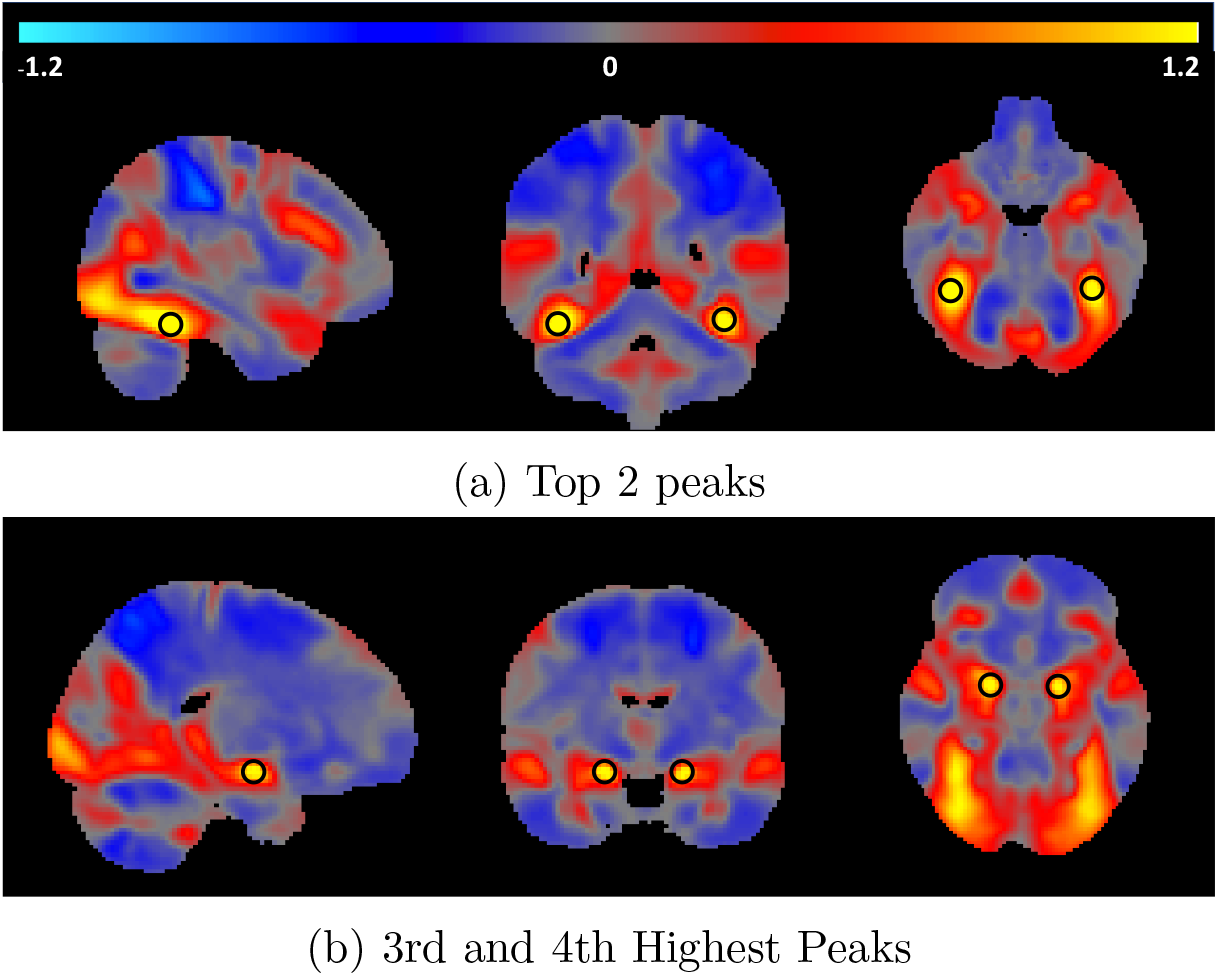
Slices through the top four maxima of the one-sample Cohen’s *d* ground truth. The top two local maxima are located at voxels (42, −46, −-22) and (−38, −48, −20) (mm MNI space) which correspond to the left and right temporal occipital fusiform cortices and have Cohen’s *d* values of 1.5756 and 1.4326 respectively. The 3rd and 4th largest local maxima are located at voxels (20,−6,−14) and (−18, −6, −14) which are within the left and right amygdalae and have Cohen’s *d* values of 1.3450 and 1.3041 respectively. The locations of these peaks are indicated using black circles.

**Figure 4:**
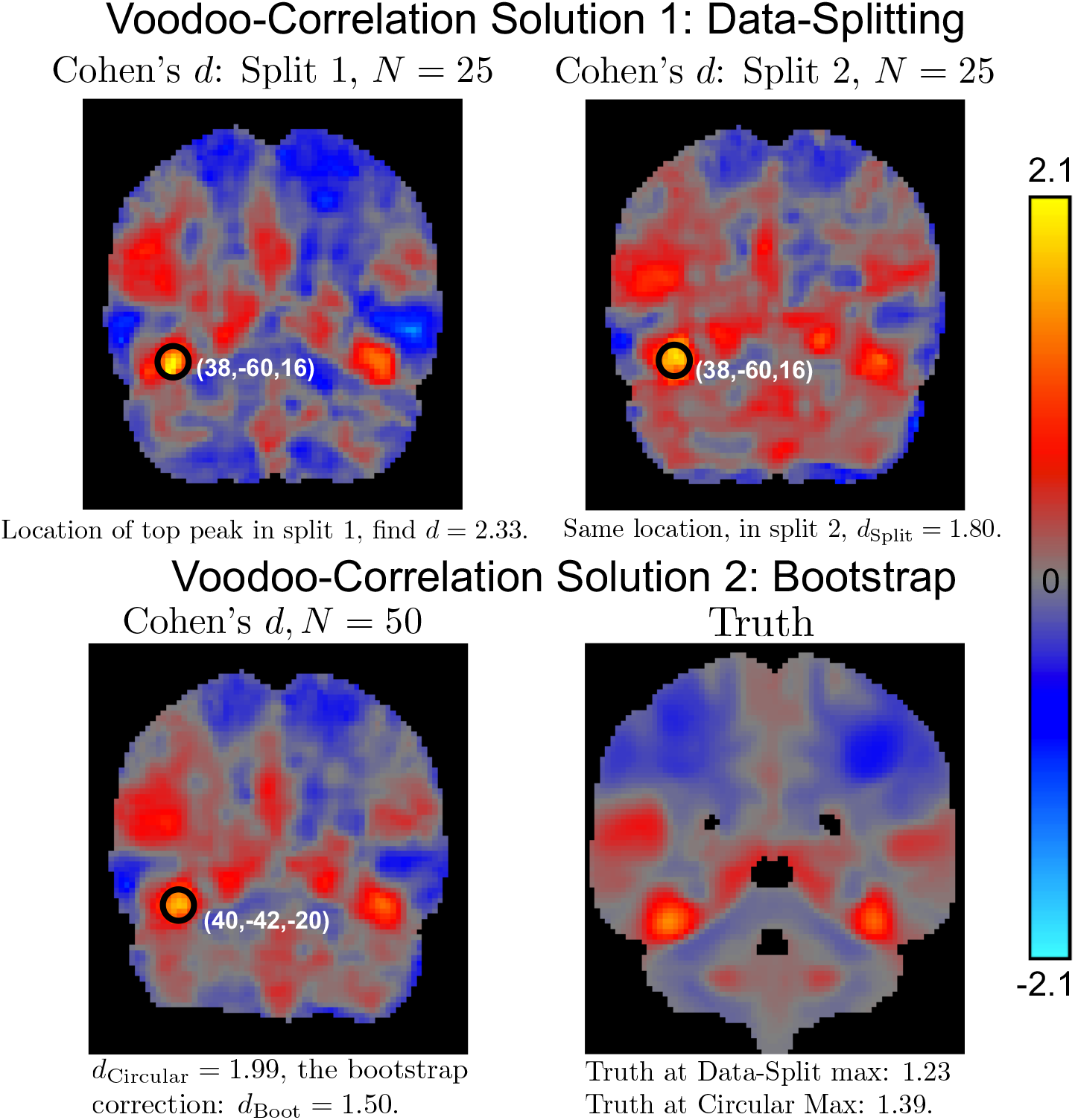
Illustration of a 50 subjects analysis using circular inference, the proposed bootstrap method and data-splitting. Data-splitting requires splitting the data in half, using the first half of the data to determine the locations, but provides a non-circular estimate. Circular inference and the bootstrap both use all of the data to calculate the locations. This figure illustrates the extra variability in data-splitting, with a noisier map and greater variability (estimated *d* of 1.80 vs truth of 1.23); the bootstrap estimate had smaller bias (estimated *d* of 1.50 vs truth of 1.39). This illustration is indicative of the extensive evaluations reported below.

#### 2.4.3 VBM data

We have structural gray matter (VBM) data from the 8,940 subjects to illustrate how our bootstrap method performs in the context of the general linear model. We regress these gray matter images against age, sex and an intercept. Using the 4,000 subjects, we calculate a ground truth estimate of the partial *R*^2^ for age at every voxel for which the mean of the VBM values at that voxel over all 4,000 subjects are greater than 0.1. The maximum of this ground truth is located at voxel (45, 62, 34) and has a partial *R*^2^ of 0.2466. As above we divide the remaining subjects into *G_N_* [4940/*N*] groups, for *N* = 50,100 and 150 to compare the methods by calculating the partial *R*^2^ for age on each subgroup. Here we use larger sample sizes as this setting is more challenging for inference as the effect size is considerably smaller than in the task fMRI data. See Section 3.2.3 for the results of this validation.

#### 2.4.4 Threshold Computation

For real data analyses, researchers typically either use random field theory (RFT) (Worsley et al., 1996) or permutation testing (Nichols and Holmes, 2002) to compute screening thresholds. Voxelwise RFT controls the false positive rates but is slightly conservative, (Eklund et al., 2016), primarily because the lattice assumption is not valid for low smoothness levels. On the other hand one-sample permutation can have slightly inflated false positive rates, (Eklund et al., 2019), and has a high computational cost. In our case this cost is very large as we need to perform a big data validation which requires many analyses as discussed in Section 2.4. We have thus elected to use voxelwise random field theory for our big data analyses. In practice when running a typical fMRI/VBM analysis, our methods work independently of the method used to choose the threshold.

### 2.5 Method Comparison

In order to compare the bootstrap, data-splitting and circular inference (in simulations and in the big data validation), we consider their bias, standard deviation and root mean squared error (RMSE) calculated over the 1,000 realizations for each *N*. Here we are computing the bias, standard deviation and RMSE in a non-standard context, in that the true parameter values vary in each instance. Traditionally in inference, one estimates a common *θ* with estimators 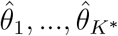, giving us the usual MSE decomposition for a sample of size *K*\*,

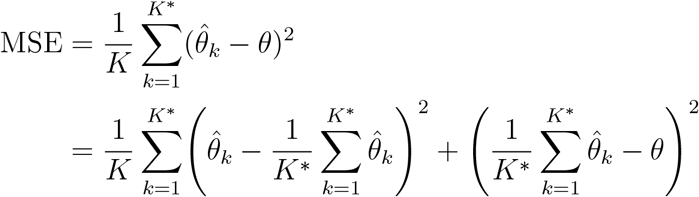

into variance and squared bias. However in our context we have estimators 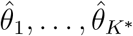 of parameters *θ*_1_, …, *θ_K_*\*. In our setting, *K*\* is the number of significant peaks that are found over all realizations. For *k* = 1, …, *K*\*, 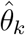 is the value of one of these significant peaks, and *θ_k_* is the true value at the voxel corresponding to the peak. The *θ_k_* are different because the locations of the peaks are random. As such we instead define

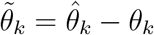

and use the fact that the noise-free value of 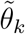 is 0 for each *k*. This allows us to define the MSE as

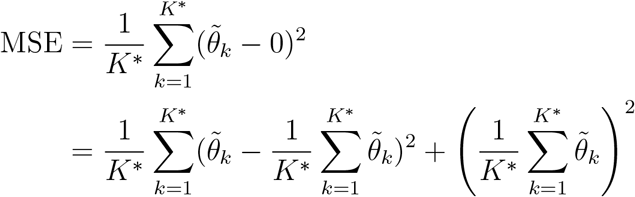

where the second equality follows by bias-variance decomposition. This leads us to define the variance

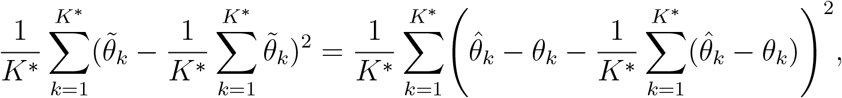

and the bias

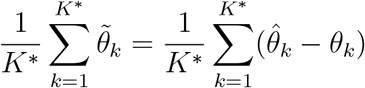

in this context. These expressions represent the bias/variance/MSE at a randomly selected peak in the sense that if a peak is selected at random we expect it to be biased above the true effect size by this amount on average. The root mean squared error or RMSE is defined to be the square root of the MSE and the standard deviation to be the square root of the variance.

In the context of the big data analysis described above, for a given sample size *N*, *K* * is the number of significant peaks over all the *G_N_* subsets. In Figures 10, 12, 14 for each set of estimates we have made boxplots for the bias 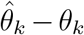 over all *K*\* significant peaks. For the bar plots in these figures we have plotted the RMSE and standard deviation as defined above.

## 3 Results

### 3.1 Results - Simulations

We found that Algorithm 1, bias correction for peak height in a sample mean image alone, had similar performance as for bias correction of statistic peaks (Algorithm 2), detailed below. However, as direct assessment of the mean image provides no way to control false positives, we regard it as a less useful method and have relegated its evaluation (on simulations and real data) to the Supplementary Material (Section S1). In this section we illustrate the performance of Algorithm 2. The performance of the methods in the GLM setting (Algorithm 3) is very similar and so this has also been left to the Supplementary Material (Section S2).

#### 3.1.1 One Sample Cohen’s *d*

Figure 5 left column plots the estimates of the bias, standard deviation and root mean squared error from each of the three methods as the sample size increases where the peak effect size is 0.5 as discussed in Section 2.3.2. As expected the circular method has the worst bias, but has low standard deviation, while data splitting is unbiased, but has the highest standard deviation. Bootstrapping has a bias which decreases to 0 as the sample size increases. Summarising mean and standard deviation with RMSE, we find that the bootstrap method has the lowest RMSE for all sample sizes except *N* = 20. The *N* = 20 exception likely occurs because resampling methods perform better for larger sample sizes.

**Figure 5:**
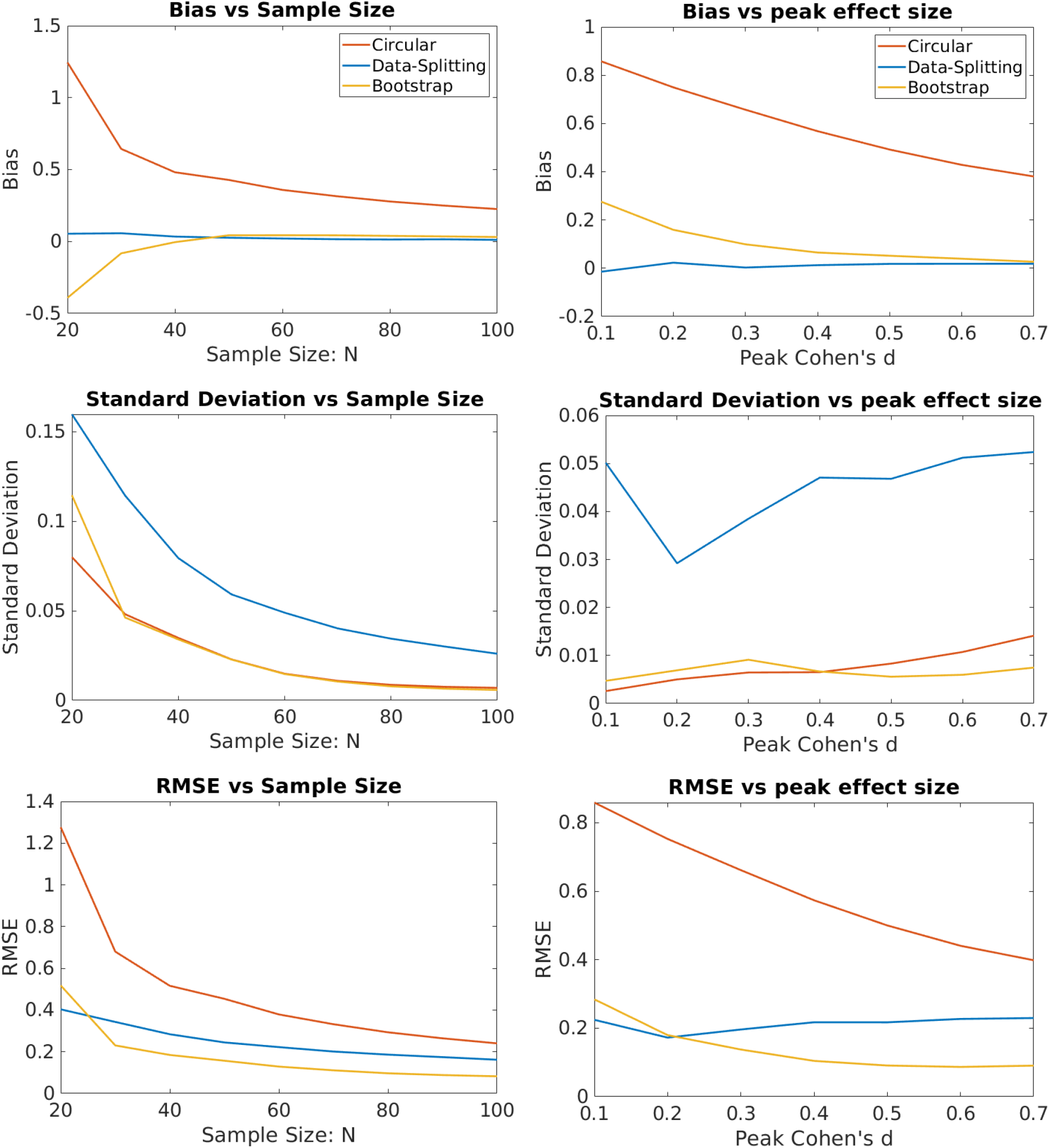
Evaluation as sample size and peak effect size changes of bias correction for Cohen’s *d* peaks (Algorithm 2) on simulated data generated as described in Section 2.3. Left column takes the effect size to be fixed and looks at how the measures change with sample size. Right column takes the number of subjects to be 50 and looks at how the measures change when the effect size is scaled. Each plot shows the bias (top), standard deviation (middle) and RMSE (bottom) calculated over 1,000 realisations. By the overall measure of RMSE, the bootstrap method performs the best except for the smallest effect sizes and sample sizes.

Figure 5 right column plots the estimates of the bias, standard deviation and root mean squared error from each of the three methods for *N* = 50 for a range of peak effect sizes. At all except the smallest effect size the bootstrap outperforms the others in terms of RMSE. The small effect size deviation occurs because the bootstrap correction is based on the rank order of the peaks, and when SNR is low the sample rank orders is a poor approximation of the noise-free rank order. As such for lower effect sizes a larger number of subjects is required for the bootstrap to outperform the other methods. For all methods the bias decreases as the peak effect size increases, this occurs because the peaks in the signal are more prominent and therefore are less subject to the winner’s curse.

In our simulations the circular and bootstrap methods find considerably more peaks than data-splitting. This is to be expected as they use double the data (relative to data-splitting) to locate the peaks and are thus more powerful. Indeed in many of our simulations for small sample sizes, data-splitting often found no peaks to be significant at all. In order to compare the power of the methods we computed the average number of significant voxels found across all realizations for each sample size (Figure 6). Circular inference and the bootstrap both use the one-sample *t*-statistic to determine significance so they both find the same number of voxels above the threshold, which is substantially more than the number found by data-splitting.

**Figure 6:**
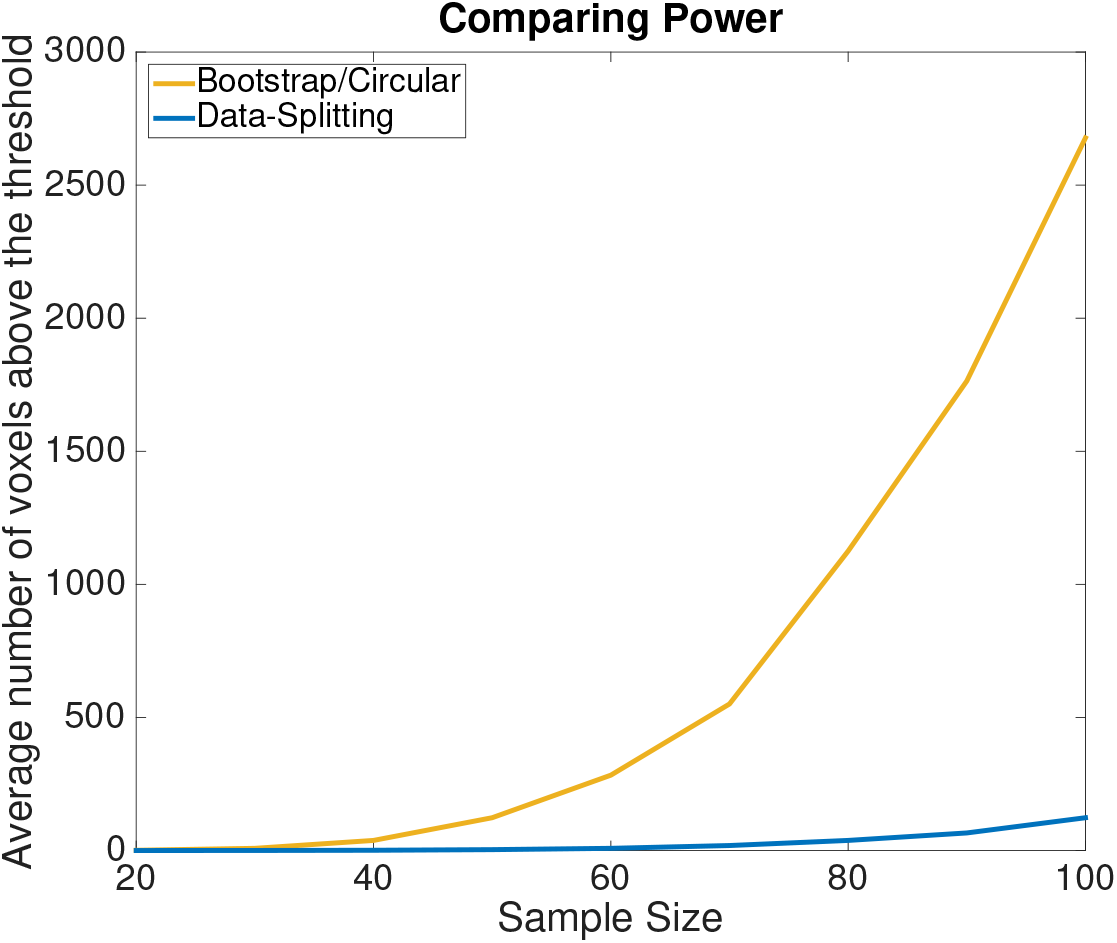
Comparing the power of the methods applied to the one-sample Cohen’s *d* simulations. We have computed the average number of significant voxels found above the threshold per realization for *N* ∈ {20, 30, …, 100} over all 1,000 realizations. Circular inference and the bootstrap find considerably more voxels to be significant than data-splitting.

#### 3.1.2 Estimating the Mean

As discussed in Section 2.1.3, a bias correction for the mean at locations of peaks in the statistic image can be obtained from a variant of Algorithm 2. Estimates of the bias, standard deviation and RMSE of each of the three methods (see Figure 7), show a very similar performance as for the previous setting, with the bootstrap method having the lowest RMSE across all sample sizes. When the SNR is low a larger number of subjects is required before the bootstrap outperforms data-splitting in terms of RMSE (see Figure 7, bottom right plot). To illustrate that this occurs we have included plots that illustrate the relative performance of the algorithms (for a larger number of subjects) in Figure S4 of the Supplementary Material.

**Figure 7:**
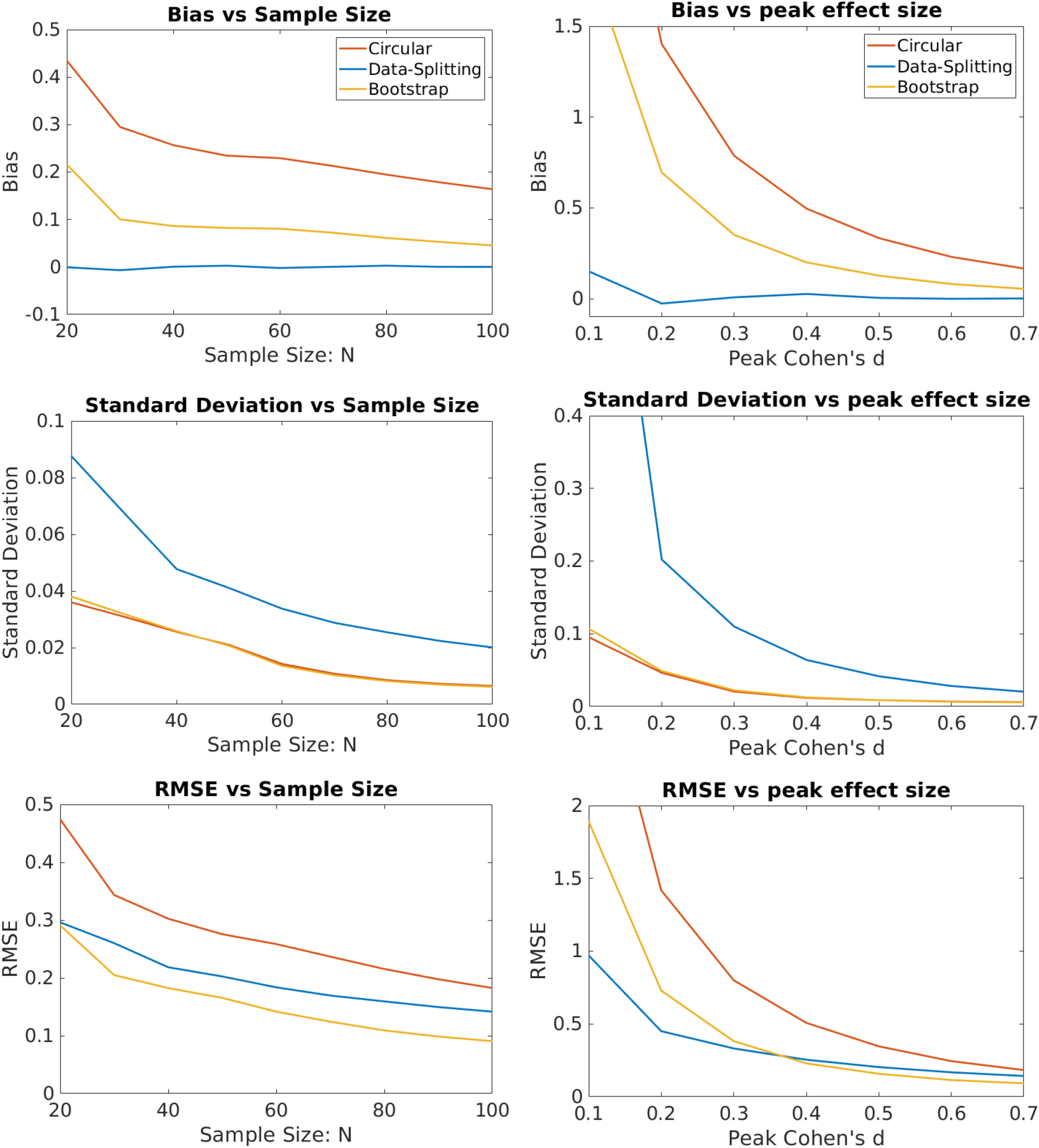
Evaluation as sample size and peak effect size changes of bias correction for the %BOLD mean at locations of Cohen’s *d* peaks (Algorithm 2) on simulated data generated as described in Section 2.3. Left column takes the effect size to be fixed and looks at how the measures change with sample size. Right column takes the number of subjects to be 50 and looks at how the measures change when the effect size is scaled. Each plot shows the bias (top), standard deviation (middle) and RMSE (bottom) calculated over 1,000 realisations. By the overall measure of RMSE, the bootstrap method performs the best except for the smallest effect sizes and sample sizes. The small effect sizes in the bottom right graph require a larger number of subjects before the bootstrap estimates attain a lower RMSE than data-splitting.

### 3.2 Results - Real Data

In this section we apply the methods to task fMRI and VBM data as described in Section 2.4. However, before discussing these these results, we illustrate the magnitude of the circularity problem we compare maximum peak heights as a function of sample size. To do so we compute the maximum peak height (of Cohen’s *d*) for different *N* ranging from 10 to 100, (averaged over the *G_N_* groups), and compare to the true max peak height of Cohen’s *d*, see Figure 8. The bias is substantial for small *N* but is non-negligible even for moderate *N*. As *N* increases the bias decreases to zero as expected and the average peak maximum converges to the true maximum value.

**Figure 8:**
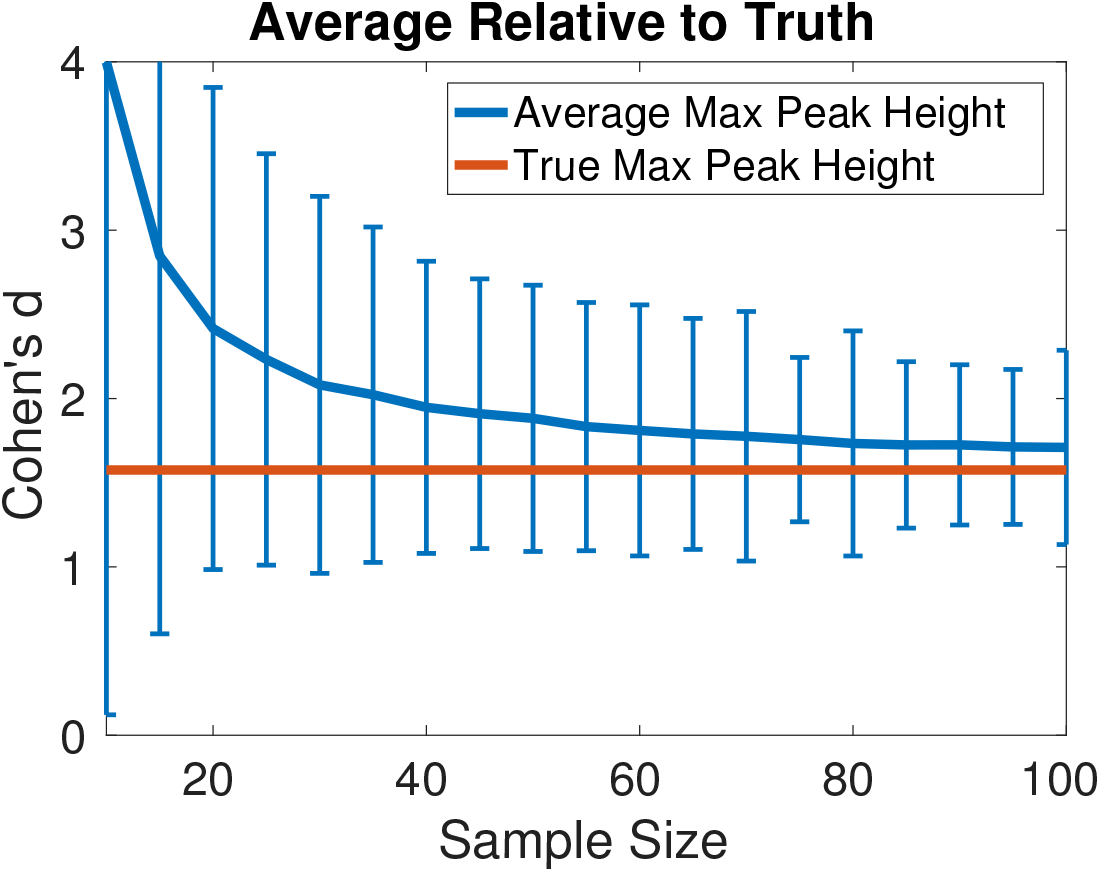
Illustration of the winner’s curse: average selection bias in the one-sample Cohen’s *d*: the average peak height of the maximum is plotted against *N*. For each *N* we compute Cohen’s *d* for each of the *G_N_* groups of size *N* find the value of the maximum and take the average over the *G_N_* groups. The 95% error bars are based on the 2.5% and 97.5% quantiles for each sample size. The bias is substantial for small *N* but is non-negligible even for moderate *N*.

#### 3.2.1 Evaluation: Task fMRI Cohen’s *d* peak height estimation

Figure 10 presents the results of applying Algorithm 2 to the one-sample task fMRI data, and is analogous to the simulated data results in Figure 5. (Note that as bias can be measured at each peak, it can be presented via boxplots; whereas only a single standard deviation and RMSE can be computed per setting, see Section 2.5 for details) As in the simulations we find that the circular estimates are highly biased whereas the bootstrap estimates have low bias with the bias decreasing as the sample size increases. The bootstrap has the lowest RMSE for each sample size.

**Figure 9:**
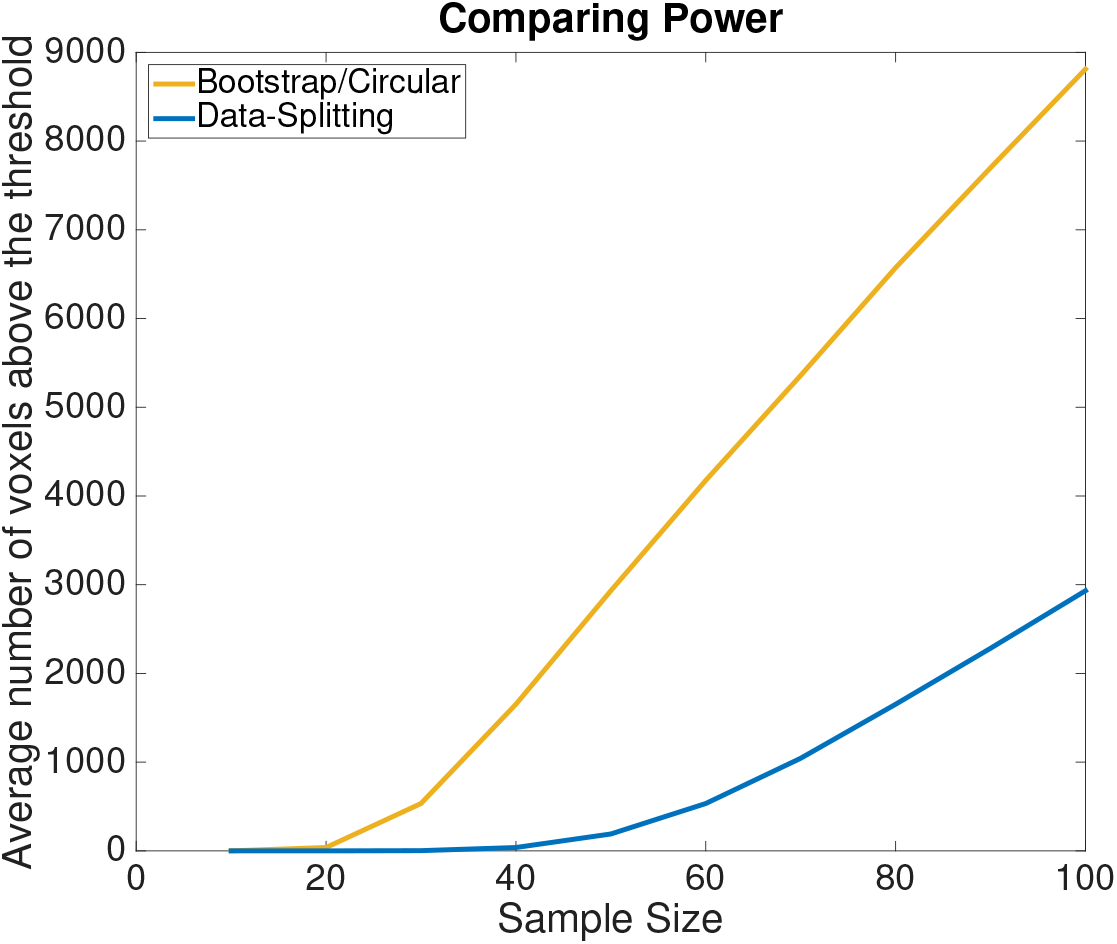
Comparing the power of the three methods. For each *N* ∈ {10, 20, …, 100} we consider the average (over the *G_N_* independent groups) number of voxels above the screening threshold. Circular inference and bootstrap find considerably more peaks than data-splitting.

**Figure 10:**
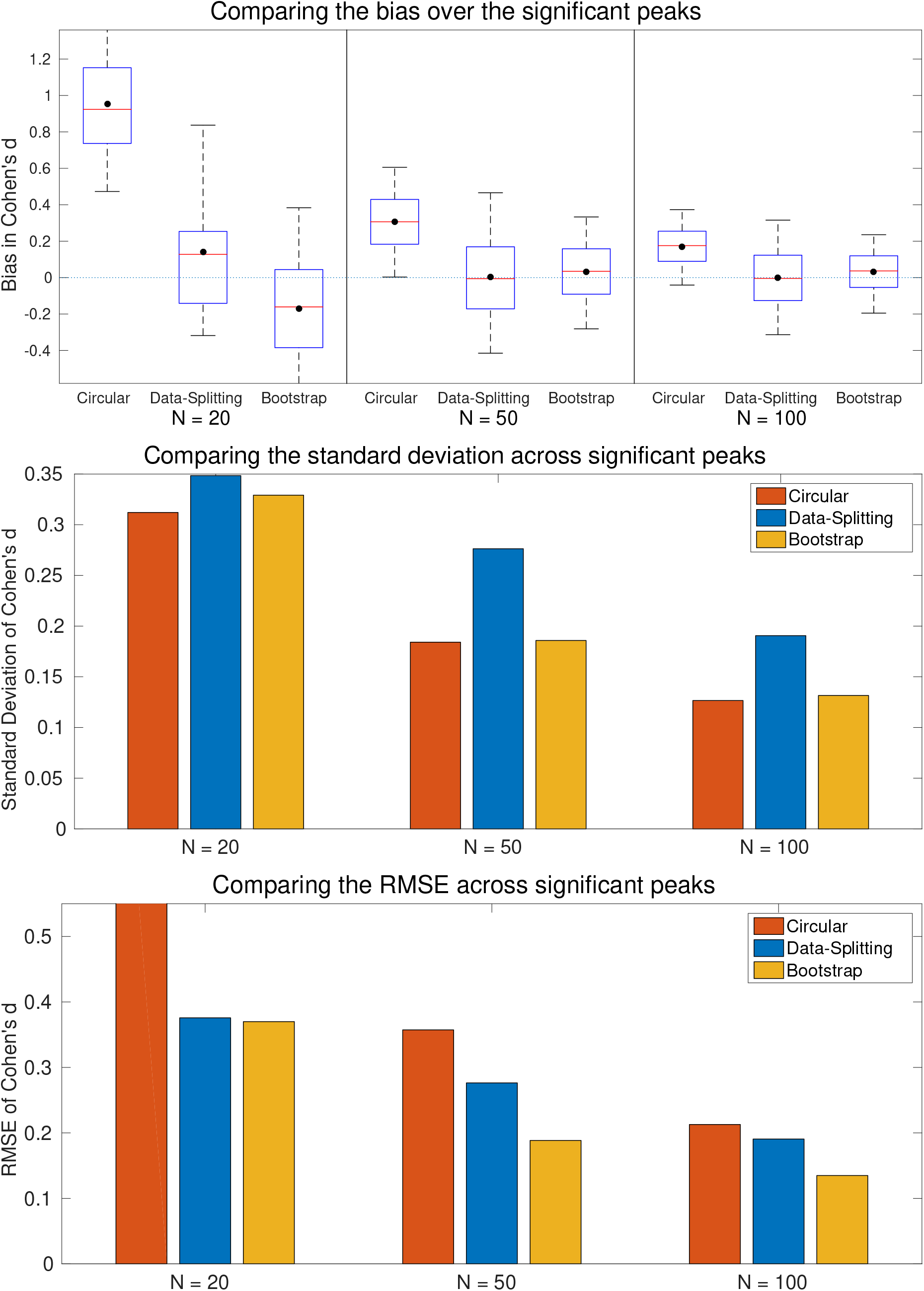
Comparison of one sample Cohen’s *d* estimation for task fMRI. Bias (top), standard deviation (middle), and RMSE (bottom) are shown for *N* = 20, 50 and 100 sample sizes, based on *G_N_* samples. While both data-splitting and bootstrap are generally unbiased, the bootstrap has the smallest RMSE. Note that the *N* = 20 data-splitting values are computed using only 7 data points so may not be representative. Note also that the RMSE for circular inference for *N* = 20 is cutoff and actually has a value of 1.0029.

In order to compare the power of the methods, as with the simulations, we have computed the average number of voxels above the threshold over all *G_N_* groups (for *N* ∈ {10, 20, …, 100}) see Figure 9. From this we see that, for a given sample size, circular inference and the bootstrap find many more peaks than data-splitting which illustrates the considerable difference in power. We note that for data splitting with *N* = 20 we observe only a total of 7 peaks that were above the threshold over all the *G*_20_ = 247 groups, so the results in Figure 10 could be unstable in this case.

To further understand how the estimates compare we plot their values against the ground truth in Figure 11. For each *N*, each data point in the corresponding graph shows the estimated peak intensity of a significant peak from one of the *G_N_* groups (ordinate), and the ground truth intensity at the location of the peak (abscissa). The *N* = 20 case is the most challenging for estimation. Here the circular estimates are very biased while bootstrap estimates give reasonable estimates and the data-splitting estimates are particularly variable and are fewer in number. As *N* increases, all of the methods perform better: the circular estimates are biased, and the data-splitting estimates are variable whereas the bootstrap estimates have low bias and variance. The effect of the threshold is particularly evident in the plots for the circular method and, to a lesser extent, for the bootstrap method.

**Figure 11:**
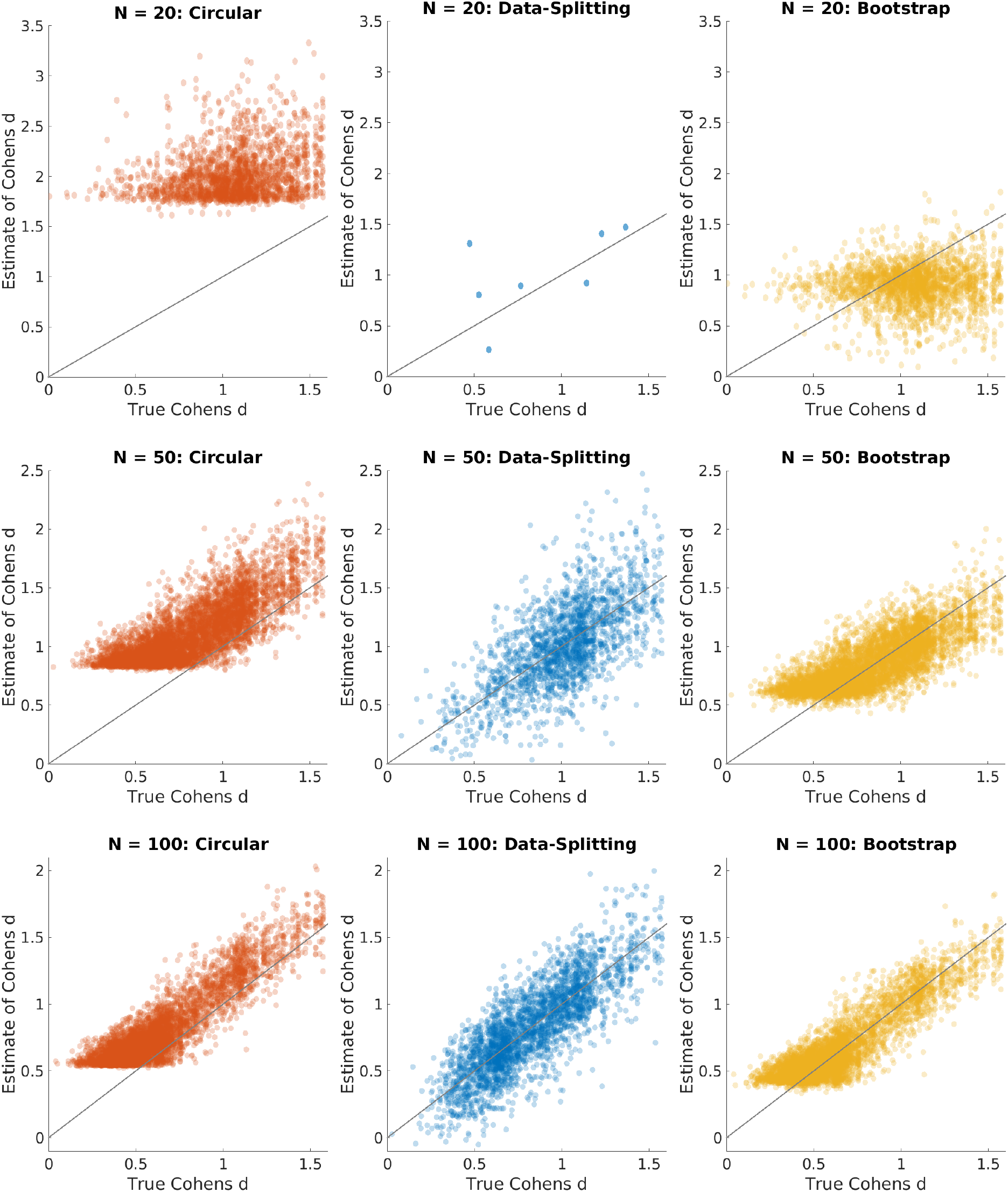
Plots of estimated versus true value of one-sample Cohen’s *d* for task fMRI images, for circular (left), data-splitting (middle), and bootstrap (right). Plots show all peaks found over the *G_N_* samples for each sample size, *N* = 20, 50,100 (top to bottom). For each peak the true Cohen’s *d* is obtained at that location from the held-out 4,000 subject Cohen’s *d* image. Note that the number of peaks and their locations are the same for circular inference and the bootstrap but are different for data-splitting as it uses the first half of the subjects in order to determine significant peaks. From these plots we can see that the bootstrap estimates have low bias and standard deviation and improve as the sample size increases. The data-splitting estimates are unbiased but are more variable and reflect fewer detected peaks.

Note that in Figure 11, for large *N*, the circular method suffers a bias that is relatively constant with respect to the true Cohen’s *d*. The bootstrap method corrects this bias, with the point cloud having roughly the same shape as the circular method’s, only shifted downward. In contrast, for the lowest *N*, there is a greater mismatch in the circular and bootstrap plots. This is because the bootstrap correction is based on the rank order of the peaks, and when SNR is low the sample rank orders is a poor approximation of the noise-free rank order.

#### 3.2.2 Evaluation: Task fMRI mean estimation at Cohen’s *d* peak location

Figure 12 illustrates the results of applying Algorithm 2 to the one-sample task fMRI data to estimate the mean at Cohen’s *d* (or *t*-statistic) peak locations. Here, all the methods perform well and have relatively low little bias, though circular inference is the most biased. Data-splitting has the worst standard deviation and RMSE for *N* = 50 and 100; it has best RMSE for *N* = 20, but we again note that this is based on only 7 peaks.

**Figure 12:**
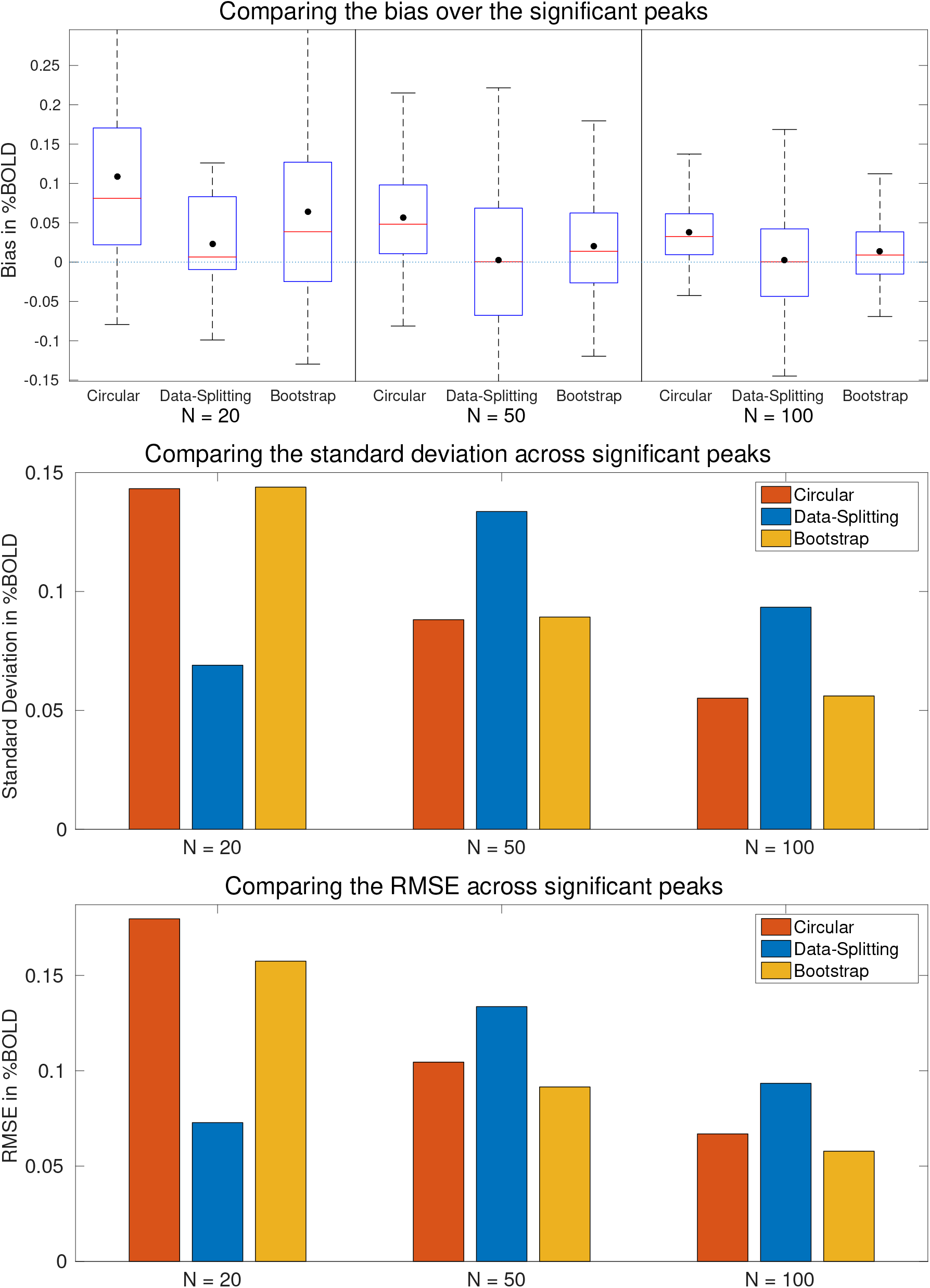
Comparison of one sample mean estimation for task fMRI. Bias (top), standard deviation (middle), and RMSE (bottom) are shown for *N* = 20, 50 and 100 sample sizes, based on *G_N_* samples. While both data-splitting and bootstrap are generally unbiased, the bootstrap has the smallest RMSE for larger sample sizes. Note that the *N* = 20 data-splitting RMSE and standard deviation is computed using only 7 data points so may not be representative. What is particular of note here is that the circular estimates have lower RMSE than data-splitting for *N* = 50 and 100.

The selection bias is much less severe in this scenario. This is due to the selection being based on Cohen’s *d*, which is correlated with but not the same as the mean. Notably for *N* = 50 and 100 the circular estimates have a lower RMSE than the data-splitting estimates, but the bootstrap estimates have the lowest RMSE.

Scatter plots comparing the estimates and the ground truth (Figure 13) reflect the observation of a much reduced problem of circularity bias; however, the bias in the circular estimates is still evident.

**Figure 13:**
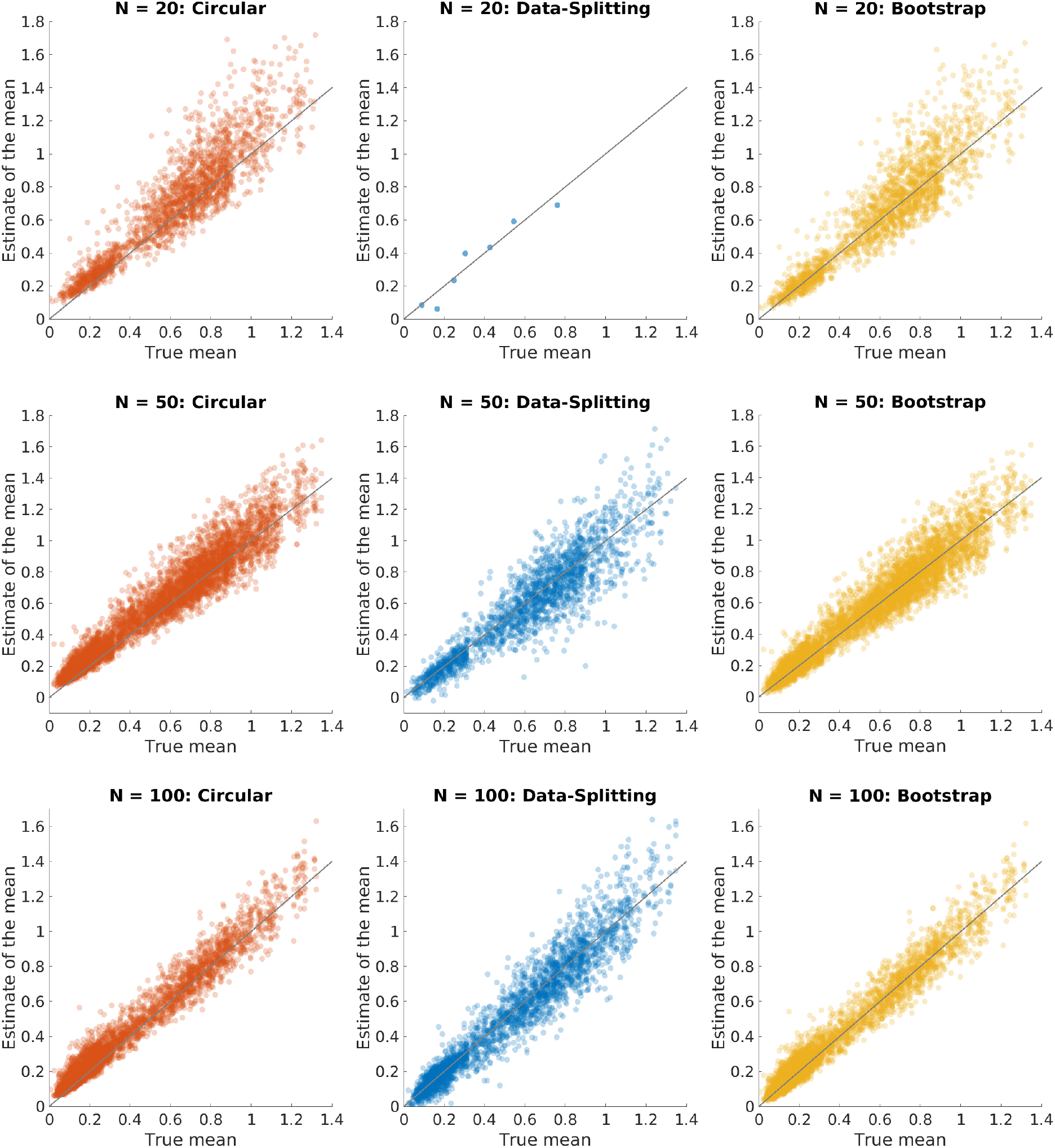
Plots of estimated versus true value of the one-sample mean (in %BOLD) for task fMRI images, for circular (left), data-splitting (middle), and bootstrap (right). Plots show all peaks found over the *G_N_* samples for each sample size, *N* = 20, 50,100 (top to bottom). For each peak the true sample mean is obtained at that location from the held-out 4,000 subject sample mean image. Note that the number of peaks and their locations are the same for circular inference and the bootstrap but are different for data-splitting as it uses the first half of the subjects in order to determine significant peaks. From these plots we see that the circularity bias is much less than for Cohen’s *d* and that the bootstrap estimates perform very well. The data-splitting estimates are unbiased but are more variable and reflect fewer detected peaks.

#### 3.2.3 Evaluation: Gray matter VBM *R*^2^ peak height estimation

Figure 14 illustrates the results for estimating the partial *R*^2^ of age with the VBM data. The performance here resembles that of Cohen’s *d* peak estimation: there is little bias for data-splitting and the bootstrap, the circular and bootstrap methods have lower standard deviation, and the bootstrap consistently has the lowest RMSE.

**Figure 14:**
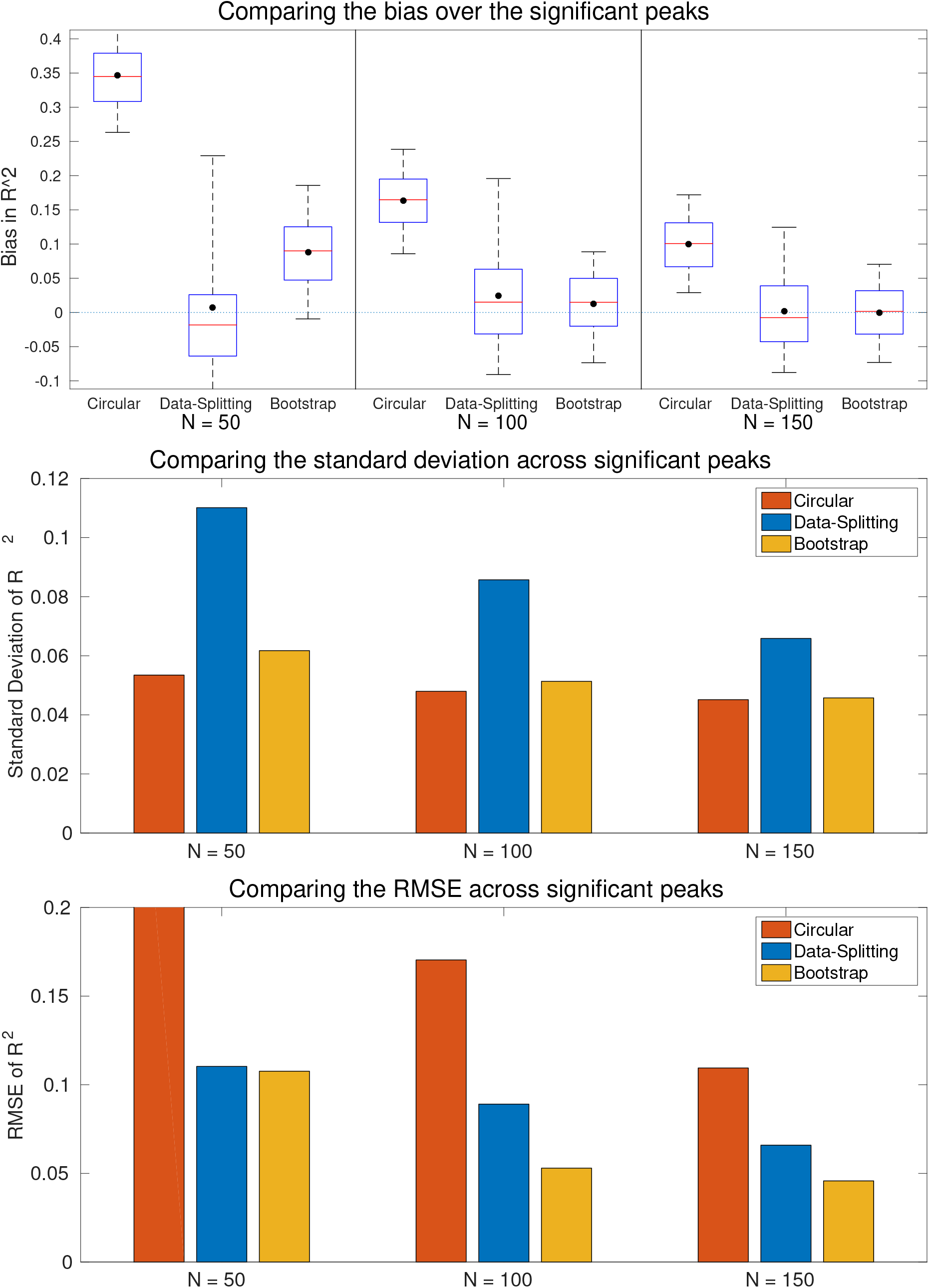
Comparison of estimates for the partial *R*^2^ for age in the presence of sex and an intercept on VBM data. Bias (top), standard deviation (middle), and RMSE (bottom) are shown for *N* = 50,100 and 150 sample sizes, based on *G_N_* samples. While both data-splitting and bootstrap are generally unbiased, the bootstrap has the smallest RMSE for all sample sizes. Note that RMSE for the circular estimates in the *N* = 50 case is 0.3407 and so is cut off by the graph.

While the boxplot and bar plot summaries (Figure 14) are consistent, the analogous Cohen’s *d* scatter plots (Figure 15) have a very different character. As the circular results (left column) make clear, most of the *R*^2^ estimates are close to the threshold, indicating a severe selection effect. As discussed above, when SNR is low the observed rank order can differ substantially from the noise-free rank order, reducing the accuracy of the bootstrap method. However, as the sample size increases the bootstrap estimates fall closer to the identity line more closely and in terms of RMSE, the bootstrap still outperforms circular inference and data-splitting.

**Figure 15:**
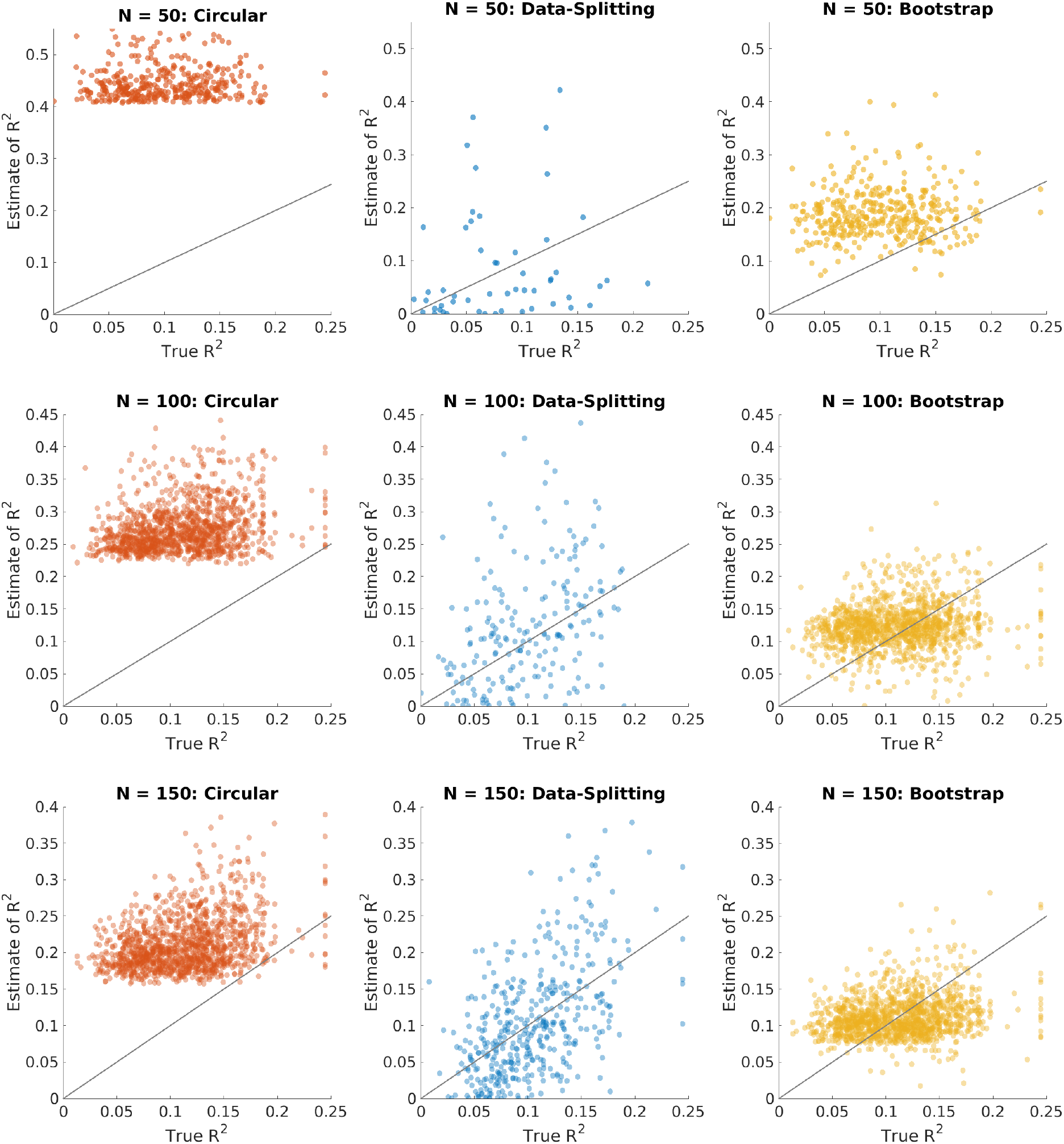
Plots of estimated versus true value of the partial *R*^2^ for age, obtained using a GLM regression on VBM data, for circular (left), data-splitting (middle), and bootstrap (right). Plots show all peaks found over the *G_N_* samples for each sample size, *N* = 50, 100, 150 (top to bottom). For each peak the true partial *R*^2^ for age is obtained at that location from the held-out 4, 000 subject partial *R*^2^ image. Note that the number of peaks and their locations are the same for circular inference and the bootstrap but are different for data-splitting as it uses the first half of the subjects in order to determine significant peaks. From these plots we see that the naive estimates are biased while the bootstrap and data-splitting estimates are unbiased on average. Data-splitting is the most variable, though the bootstrap over corrects values with a large true partial *R*^2^ and under corrects those with a low partial *R*^2^ for *N* = 100, 150. On average the bootstrap estimates lie closer to the identity line than the data-splitting estimates resulting in the decrease in RMSE, see Figure 14.

### 3.3 Demonstration on HCP Task fMRI dataset

In order to illustrate the bootstrap method in action we apply it to a sample of 80 unrelated subjects from the Human Connectome project and look at one of the working memory contrasts. Subjects performed an *N*-back task using alternating blocks of 0-back and 2-back conditions with faces, non-living man-made objects, animals, body parts, house and words. We examine the average (2-back – 0-back) contrast, identifying brain regions supporting working memory in general.

We use a group level model and compute a one-sample *t*-statistic at each voxel in order to test for activation. Voxelwise permutation testing is used to control the familywise error rate to 5% resulting in a threshold of 5.10 for the *t*-statistic. The largest peak above this threshold has a *t*-statistic value of 13.58 and lies within the Medial Frontal Gyrus an area commonly associated with working memory. At the largest peak the circular Cohen’s *d* is 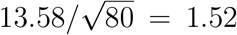; the bootstrap corrected Cohen’s *d* estimate is 1.161. In total 234 peaks lie above the threshold, with 25 peaks falling within the Medial Frontal Gyrus region (Harvard-Oxford Atlas). Table 2 reports the circular and bootstrapped Cohen’s *d* as well as the bootstrap estimate of the mean for the top 10 of these 25 (Cohen’s *d*/*t*-statistic) peaks. Slices through the one-sample *t*-statistic at the voxel corresponding to the largest peak are shown in Figure 16.

**Table 2:**
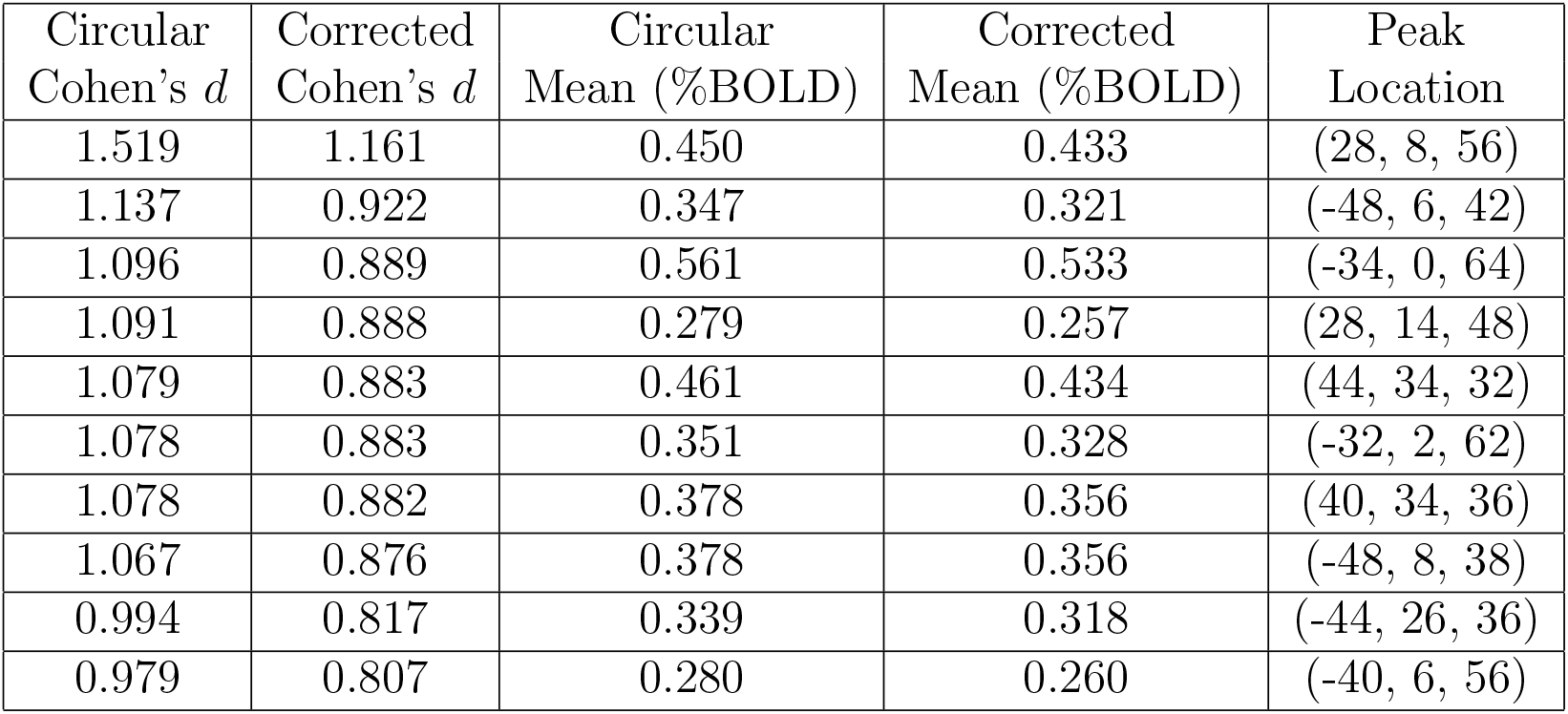
The circular and corrected estimates of Cohen’s *d* and the mean at the top ten significant peaks of Cohen’s *d* in the Medial Frontal Gyrus. There is appreciable bias for the largest Cohen’s *d* peaks, while the %BOLD values at the Cohen’s *d* peaks have relatively little bias.

**Figure 16:**
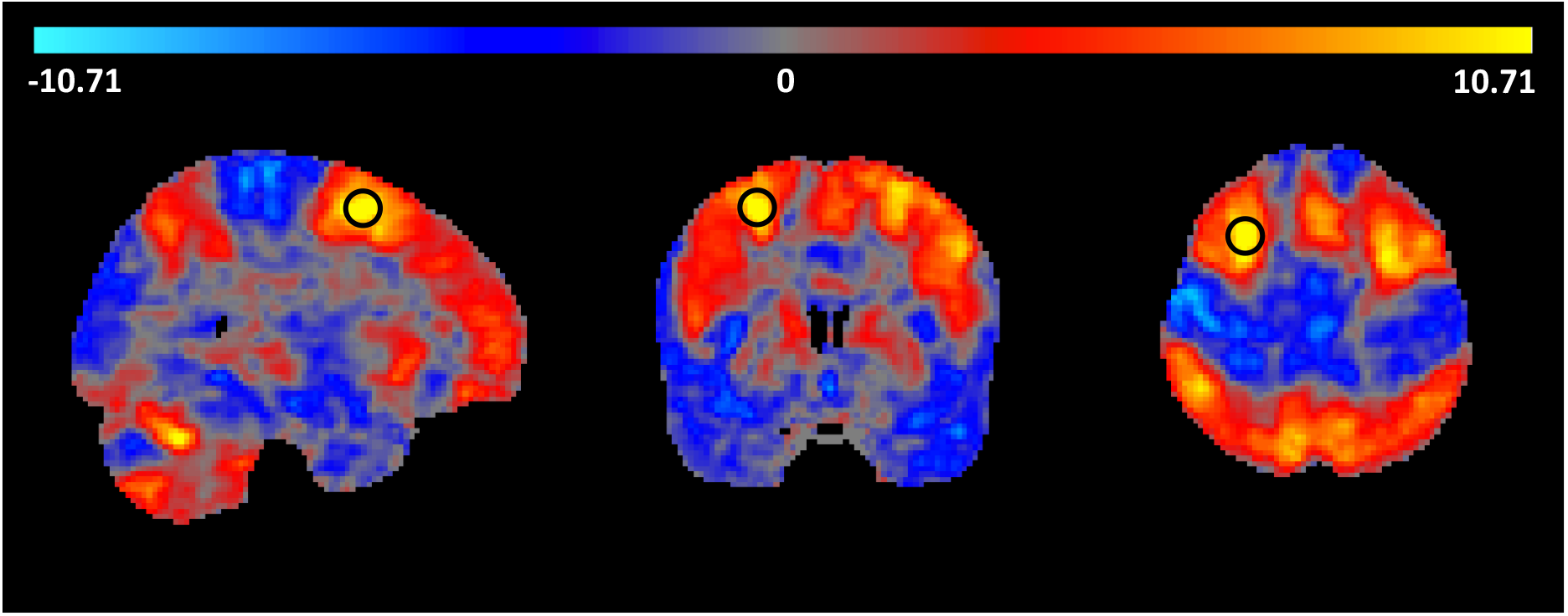
Slices through the one-sample *t*-statistic for the working memory contrast (2-back – 0-back) for subjects from the Human Connectome Project. Black circles indicate the location of the largest peak of activation which lies at the voxel (28, 8, 56) at the edge of the Medial Frontal Gyrus. At this location the observed (circular) Cohen’s *d* is 1.519, while the bootstrap-corrected value is 1.161; the observed %BOLD change at this voxel is %0.450 and corrected estimate is %0.433.

Figure 17 shows the effect that these corrections have on power, where we have plotted a graph of sample size against power for a whole brain analysis using a *p*-value threshold, corresponding to taking *T* = 5.10, of 1.39 × 10^−6^. Using the raw value would suggest that only 24 subjects are need to attain 80% power, when in fact the corrected estimate shows that 34 subjects are needed to provide this level of power. See Appendix D.2 for details on how the power is calculated.

**Figure 17:**
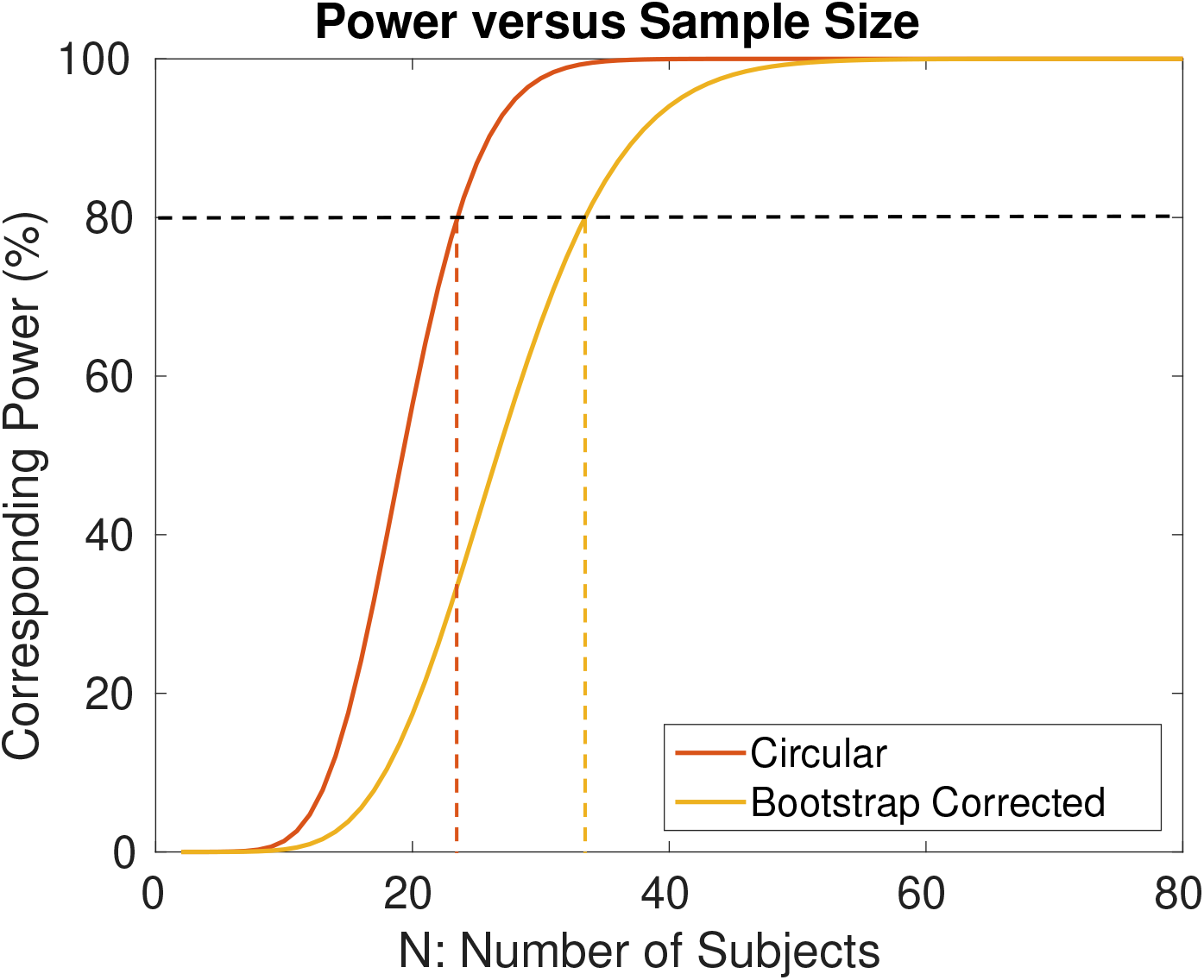
The power corresponding to a given sample size for the one-sample *t*-statistic when the *T*-statistic threshold is 5.10. The blue curve is the power curve corresponding to the circular estimate of the Cohen’s *d* for the HCP working memory dataset and the red curve is the power curve corresponding to the corrected Cohen’s *d*. Using the raw value would suggest that only 24 subjects are need to attain 80% power, when in fact the corrected estimate shows that 34 subjects are needed to provide this level of power.

## 4 Discussion

Unbiased estimation of effect size is essential yet absent from most neuroimaging studies. We have evaluated three methods for assessing the signal magnitude at peaks in neuroimaging analyses. The bootstrap method that we have introduced provides circularity-corrected estimates from an analysis using all of the data. Compared to uncorrected, circular inference our method has dramatically less bias and lower RMSE. While data-splitting is unbiased by construction, our method has lower standard deviation and RMSE in most settings. Given the small size of many studies, using data-splitting, and thereby having to divide the data in half, may produce unacceptable reductions in power.

Even for small sample sizes the bootstrap has similar or better RMSE relative to data-splitting. However, we note that in neuroimaging it is very important to have an accurate estimate of the location of the effect. For this reason we assert, that even in this scenario, the bootstrap is to be preferred over data-splitting since it uses all data to compute the peak locations. It thus identifies a greater number of significant peaks and its estimates of their locations are more accurate.

The dramatic difference in the plots comparing the estimates with the ground truth for Cohen’s *d* and *R*^2^ (Figures 11 & 15, respectively) should be viewed in terms of the dramatic difference in power between these two settings. Consider that, in these evaluations, we have a typical Cohen’s *d* of 1.0 and *R*^2^ of 0.1. If we consider a power analysis with a whole brain *α* = 1.39 × 10^−6^ (here we use the *α* level of Section 3.3 as representative of typical threshold) and a target power of 1 – *β* = 80%, a one-sample *t*-test with this Cohen’s *d* would require 42 subjects; in contrast, a simple linear regression with this *R*^2^ would require 306 subjects. In this light, we find the *R*^2^ results even more impressive, providing adequate performance even with negligible power.

Large scale repositories of neuroimaging data have enabled us to validate our methods in a way that has (to our knowledge) not been done before in the neuroimaging setting. This involves setting aside a large number of subjects to compute an accurate version of the truth and dividing the remaining subjects into small groups on which to test the performance of methods relative to the ground truth. This approach enables methods to be rigorously tested and ensures that they work on real data rather than just on simplified simulations. We recommend this sort of evaluation for all new statistical imaging methods, as well as for existing methods that have not been rigorously tested on real data.

At present our method provides a bias correction for the intensity at the location of the observed peaks 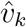. This is appropriate because when a researcher comes to replicate the results they should be able to test the effect at a given location. However another direction would be to obtain estimates for the signal intensity at true peak locations. Let v_k_ be the location of the *k*th largest peak in the noise-free image. In the setting of Algorithm 1, at present we infer on 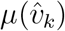 by estimating the bias 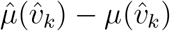, but we could instead infer on *μ*(*v_k_*) by estimating the bias 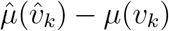. It may be possible to estimate this bias using a bootstrap approach: comparing bootstrapped peaks to peaks of the empirical mean. Peaks could be matched according to their order statistics or to the nearest large empirical peak within a certain radius in order to obtain an estimate of the bias. The challenge of this approach would be to obtain a good criteria for peak matching. Algorithms 2 and 3 could be extended similarly.

There is much ongoing research in the field of selective inference and there is much potential for other methods to be modified for use in the fMRI setting. It would particularly desirable to derive theoretical corrections using random field theory, however it is at the moment difficult to estimate the peak height distribution of a non mean zero random process (Cheng and Schwartzman, 2015), so this is an important area for future research.

Clusterwise inference is commonly used in fMRI and it is also of interest to develop selective inference approaches that allow for power analyses in this context. One approach would be to obtain an unbiased estimate of clusterwise mean (which typically suffers from selection bias) and to report the mask of where the activity lies. Our method cannot be directly used to estimate of the cluster mean because of a lack of pivotality however it may be possible to modify it in such a way that this it not a problem. Methods such as that developed in Hayasaka et al. (2007) could then be used to perform power analyses. This would provide an approximate estimate of the power though two potential issues with this approach are that in reality the mean is not constant over each cluster and that not every voxel within a significantly cluster is active.

We have motivated peak-level inference for its use in power analysis, however it also forms an essential part of how results are presented in SPM. Ever since a revision of SPM5 that introduced FDR inference for peaks, the “voxel-level” column label has been replaced with with “peak-level” in the inference table. (Confusingly, FWE *p*-values are identical for voxels and peaks, while FDR *p*-values differ substantially, with peak *p*-values notably depending on a screening threshold). Chumbley et al. (2010) have stridently argued against voxel-level inference, asserting that only peaks (and clusters) should be objects of inference in neuroimaging, as these are topological characteristics that can be unambiguously identified in a continuous process analogue of the statistic image. In general we see the value of voxel, peak and clusterwise inference, however the ubiquity of reporting peaks in statistic images in SPM and other packages is an important motivation for this work.

In sum the bootstrap approach provides a method to remove the bias while using all of the data to obtain accurate estimates of the locations. Relative to data-splitting and circular inference this results in estimates which have similar or generally better RMSE.

## 5 Software Availability and Reproducibility

The analysis in this paper was performed using MATLAB 2015a. Scripts to implement the bootstrap, circular inference and data-splitting methods are available at https://github.com/sjdavenport/SIbootstrap. Code to perform power analyses and large-scale linear modeling has also been included. For reproducibility scripts to reproduce the figures in the results section are also available in the Results_Figures folder.

Simulations and thresholding were performed using code from the RFTtoolbox available at https://github.com/sjdavenport/RFTtoolbox. Brain imaging figures where created using FSLeyes (McCarthy, 2019).

## 6 Acknowledgments

We would like to thank the 3 anonymous reviewers for their comments which have helped to improve the quality of this manuscript.

TEN is supported by the Wellcome Trust, 100309/Z/12/Z and SJD is funded by the EPSRC. Data were provided in part by the Human Connectome Project, WU-Minn Consortium (Principal Investigators: David Van Essen and Kamil Ugurbil; 1U54MH091657) funded by the 16 NIH Institutes and Centers that support the NIH Blueprint for Neuroscience Research; and by the McDonnell Center for Systems Neuroscience at Washington University.

## Appendices

### A Computing partial *R*^2^ from an *F*-statistic

For a general linear model, let Ω denote the overall model and let *ω* ⊂ Ω denote some sub-model with *p*_0_ degrees of freedom. Define RSS_⊓_ and RSS_*ω*_ to be the residual sum of squares for each of the models. Then we can write the *F* statistic for comparing *ω* and Ω as

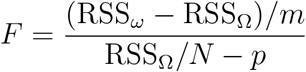

where *m* = *p* – *p*_0_ and the partial coefficient of determination is:

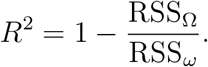

Thus, with some algebra *F* can be expressed in terms of *R*^2^ as

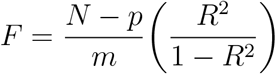

and conversely, *R*^2^ can be expressed in terms of *F* as,

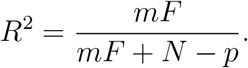

The *F*-statistic above has a different form to the *F*-statistic defined in Section 2.2. For every *m* × *p* contrast matrix *C* taking the sub-model *ω_C_* = {*β* : *Cβ* = 0} and applying the General Linear Hypothesis establishes their equivalence.

### B Masking and Calculating the Ground Truth

The UK Biobank enables us to set aside a large number of subjects in order to get a very accurate estimate of the true effect size, be it the mean, Cohen’s *d* or a regression coefficient or partial *R*^2^ in a linear model. This appendix describes the details of how we use such a large sample to create ground truth and how we deal with practical challenges, including masking and the inability to load all images into memory at once.

#### B.1 Masking

Any neuroimaging analysis requires a mask to define voxels that are to be included in the modeling process. In practice, the mask for each subject is unique. Let 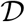 be the set of all possible voxels in the image, then given a subject: *n*, define its **mask** to be the image: 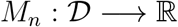 such that *M_n_*(*v*) is 1 if subject *n* has data at voxel *v*, 0 otherwise. Given this definition, define the **intersection mask** of a subset 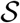 of subjects to be the image 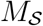 such that

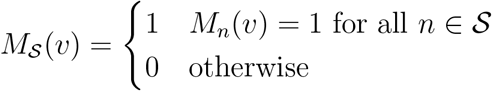

The mask used for a small sample analysis on the subset 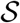 is the product of the intersection mask 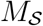 with the 2mm MNI brain mask, the image MNI152_T1_2mm_brain_mask in FSL. We refer to this as the **analysis mask**.

#### B.2 One-Sample Ground Truth Mean and Cohen’s *d*

We choose a random subset 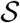 of {1, …, 8940} of size 4,000 to estimate a ground truth mean and Cohen’s *d* using the the available data at each voxel. Note that 99.99% of voxels had data from from at least 100 subjects, while 98% had data from at least 3,000 subjects. In order that each voxel be a reliable estimate we require that at least 100 subjects have data at that voxel in order that it be included. Given subject images 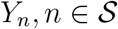, define the **ground truth mean** to be

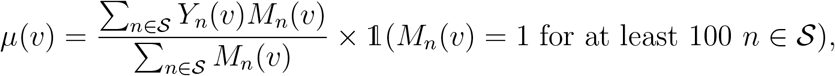

where 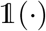 is the indicator function. Define the **ground truth variance** to be:

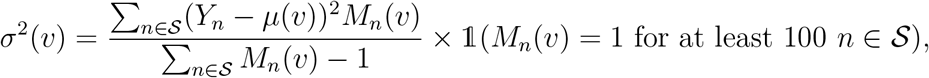

and the **ground truth Cohen’s** *d* estimate as

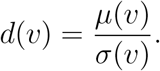

Finally each of these are additionally masked with the 2mm MNI brain mask.

#### B.3 Linear Model for Big Data - No Missingness

The full unmasked images comprise 902, 629 = 91 × 109 × 91 voxels and for 4,000 subjects this data would occupy 27GB RAM at double precision, presenting serious computational challenges. Here we outline a method for computing linear models when the data cannot be loaded into RAM all at once. Fitting separate linear models at each voxel sequentially is slow as it requires access to all of the images for each of the voxels (in the subset of the brain that is of interest) in turn. Loading all of the images at once is generally not feasible due to memory constraints. An improvement would be to divide the brain image into blocks that can fit in memory, however this still requires each image be accessed multiple times. Instead it is possible to write the estimates of the linear model in terms of individual contributions of each subject, allowing arbitrarily large datasets by only reading one subject’s data at a time. Suppose that we have *N*_all_ subjects (when computing the ground truth *N*_all_ = 4,000), that we have an *N*_all_ × *p* design matrix *X* and that there is no missing data. Let *Y* be the *N*_all_ × *V* matrix of all the subject images where *V* is the number of voxels in each subject image *Y_n_*. For the mass univariate linear model *Y* = *Xβ* + *ϵ*, we want to compute

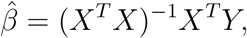

at each voxel. Instead of computing this directly we observe that for each 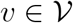,

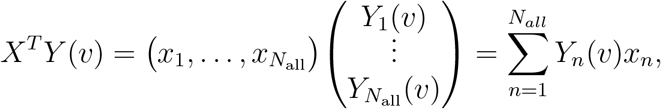

where 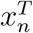 is the *n*th row of *X*, and so *X^T^Y* can computed by loading one image at a time. This *p* × *V* matrix can then be pre-multiplied by (*X^T^X*)^−1^, which only has to be calculated once, in order to calculate 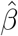. The sample variance image can then be computed by a second pass through the data as

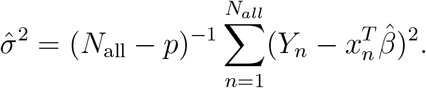

The *F*-statistic can then be computed as usual and this allows calculation of the ground truth partial *R*^2^ using the transformation from Appendix A.

#### B.4 Linear Model for Big Data - Accounting for Missingness

The previous section assumed identical masks for all subjects, which is not realistic due to susceptibility drop-out in fMRI, variation in field of view in structural MRI, and simply random variation in the exact brain boundary in each subject. For each subject *n* = 1, …, *N*_all_, suppose that we have a binary mask image *M_n_* which denotes the missingness in the response (we will assume that there is no missingness in the predictors). Since there is no missingness in the covariates and they are fixed, we can obtain an unbiased estimator of the regression coefficient at each voxel using the complete data at that voxel, so long as we assume that the missingness mechanism is independent of the image data (White and Carlin, 2010). Assuming this is reasonable, given that the missingness is typically due to acquisition and technical artifacts.

For each voxel *v*, let *C*(*v*) := {*n* : *M_n_*(*v*) = 1}, and let *C*(*v*) as a subscript indicate subsetting the corresponding rows of a matrix. Then for each voxel *v* we need to compute

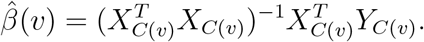

The first and second parts of this expression can be computed as

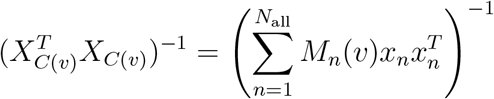

and

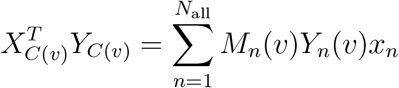

so this can also be computed by loading one image at a time. This requires storage of a *p* × *p* matrix at each voxel which, for moderate *p*, is not problematic. The same masking computation can be used when computing residual variance, which then allows computation of the *F*-statistic and this allows calculation of the ground truth partial *R*^2^ using the transformation from Appendix A.

### C Neighbourhoods and Local Maxima

Suppose that the vertices in 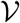 are connected by a set of edges. Let the collection of these edges be denoted by *E*. Then we define two vertices *u* and *v* to be **neighbours** in the graph 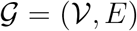 if the edge connecting *u* and *v* is contained in the set of edges **E**. Given 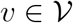, define the **neighbourhood** of *v* to be the set of voxels that are neighbours to *v* and denote this by *ne*(*v*).

Given an image 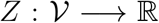, we define a voxel *v* to be a **local maxima** if *Z*(*v*) ≥ *Z*(*v*′) for all *v*′ ∈ *ne*(*v*). In 3D brain images we have a rectilinear grid of voxels and take the edge set to be defined by a connectivity criterion of either 6, 18 or 26, which if our voxels are represented by cubes correspond to those surrounding voxels which share surfaces, edges and corners respectively. As such which voxels are defined to be local maxima is dependent on the connectivity criterion. In this paper we have used a neighbourhood criterion of 18 which is the default in SPM.

### D Non-Central Distributions and Power Analyses

#### D.1 Non-Central Distributions

##### D.1.1 One-Sample *t*-statistic

Following the model from Section 2.1, under the assumption of Gaussian noise,

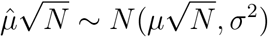

is independent of 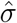 and so the *t*-statistic 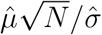 has a non-central *t*-distribution with non-centrality parameter 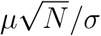 and *N* – 1 degrees of freedom. The mean of the noncentral t is not the non-centrality parameter, instead

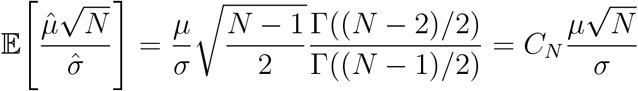

for *N* > 2, where Γ is the gamma function and *C_N_* is a bias correction factor (Hogben et al., 1961). Thus we use

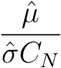

as an unbiased of the population Cohen’s *d*. Note that here and henceforth whenever we have two images *A* and *B* we write 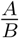 to be the image which takes the values 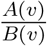 at each voxel *v*.

##### D.1.2 Non-Central *F* and *t* distributions in the General Linear Model

For the general linear model at a given voxel (note we suppress the voxel *v* index here), we have 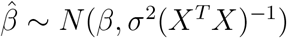 independently of 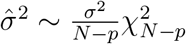 which implies that

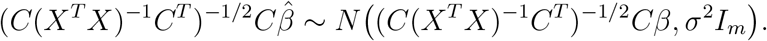

Thus 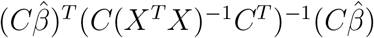 has a non-central chi-squared distribution with *m* degrees of freedom and non-centrality parameter (*Cβ*)^*T*^(*C*(*X^T^X*)^−1^*C^T^*)^−1^(*Cβ*). In particular

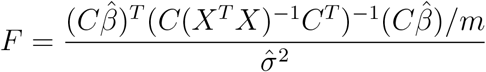

has a non-central *F* distribution with non-centrality parameter

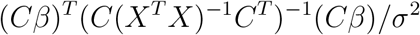

and degrees of freedom *m* and *N* – *p* and so

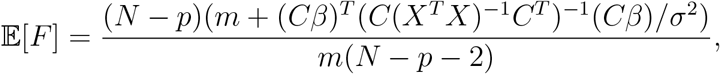

as derived in Patnaik (1949). In the case where *C* = *c^T^* is just a single contrast vector and we want to perform inference using the *t*-statistic instead of the *F*-statistic, the *t*-statistic

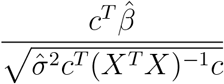

has a non-central *t*-distribution with *N* – *p* degrees of freedom and non-centrality parameter 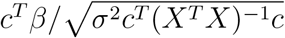.

#### D.2 Power Analyses

##### D.2.1 One Sample

In the one sample scenario, for a potential future sample size *N*′ and an estimate of the non-centrality parameter: λ, the power is:

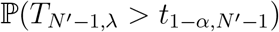

where *t*_1–*α,N*′–1_ is chosen such that 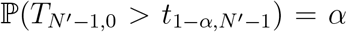 and *T*_*N*′–1,λ_ has a noncentral *T* distribution with *N*′ – 1 degrees of freedom and non-centrality parameter λ.

##### D.2.2 Multiple Regression - Cohen’s *f*^2^

Calculation of power in the general linear model scenario is slightly more complicated as it requires distribution assumptions and approximations. To do so define Cohen’s *f*^2^ to be

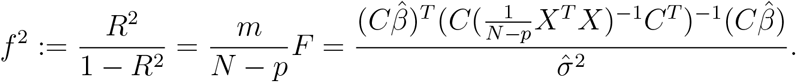

where *R*^2^ is the partial coefficient of determination and we have used the fact that 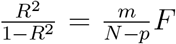 as derived in Appendix A. In the framework of the general linear model (2), for 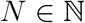 suppose we observe an N-dimensional image *Y_N_* such that

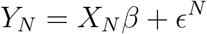

for some *p*-dimensional parameter image *β* and *N*-dimensional noise image *ϵ^N^* = (*ϵ*_1_, …, *ϵ_N_*)^*T*^ where {*ϵ_n_*}_*n*∈*N*_ is an iid sequence of noise images which have finite variance. Let *X_N_* = [*x*_1_, …, *x_N_*]^*T*^ be the design matrix, where 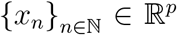 is a sequence of finite variance, iid random vectors (independent of the noise process) each with multivariate distribution *D*. For each *N*, let 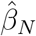 be the *p*-dimensional image linear least squares estimator and let 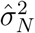 be the image estimate of variance.

Then 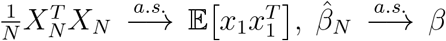 and 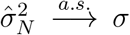 as *N* → ∞ (where 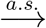 denotes pointwise almost sure convergence) see the supplementary material Section S6 for proofs. Let 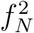 be Cohen’s *f*^2^ for the *N*th model. Then combining the above results,

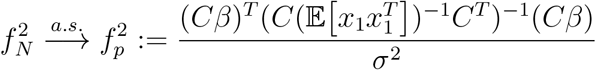

as *N* → ∞. This also implies almost sure convergence of *R*^2^.

Given a new sample of *N*′ subjects from model D.2.2 with corresponding design matrix *X*′ (an *N*′ × *p* matrix whose rows are iid with distribution *D*), then as long as *N*′ is sufficiently large, we can obtain reasonable estimates of the power. To do so note that the (new) *F*-statistic has a non-central *F* distribution with non-centrality parameter:

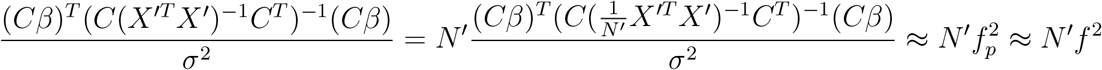

where *f*^2^ is the estimate of 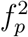. Let λ = *N*′*f*^2^ be the estimate of the non-centrality parameter. Then the power is:

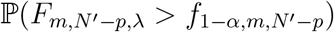

where *f*_1–*α,m,N*′–*p*_ is chosen such that 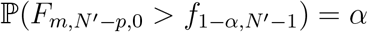 and where *F*_*m,N*′–*p*,λ_ has a non-central *F* distribution with *m* and *N*′ – *p* degrees of freedom and non-centrality parameter λ.

##### D.2.3 Multiple Regression - Cohen’s *f*

In the case that *C* = *c^T^* is a contrast vector, we often use the *t*-statistic as this allows us to perform one-sided tests. In which case we can use Cohen’s *f* which is defined as

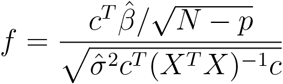

and use 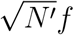 as our estimate of the non-centrality parameter using this to calculate an estimate of the power.

## Footnotes

1 In voxelwise inference voxels with test statistic values lying above a multiple testing threshold are determined to be significant. In both peak and cluster level inference a primary threshold is used to identify peaks/clusters and then thresholding based on peak magnitude and cluster extent is used to determine significant peaks/clusters.

2 https://biobank.ctsu.ox.ac.uk/crystal/docs/brain_mri.pdf

3 For 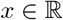, [*x*] is the largest integer that is less than or equal to *x*.

