## Supplementary Material for "Selective peak inference: Unbiased estimation of raw and standardized effect size at local maxima"

September 5, 2019

#### Contents

**S1 Application of Algorithm 1 to Simulated Data** **S1**

**S2 GLM simulations** **S1**

**S3 Additional Simulations for Estimating the Mean at Cohen's $d$ peaks** **S3**

**S4 Application of Algorithm 1 to fMRI Data** **S3**

**S5 Comparing the Bootstrap and Circular Inference at top peaks** **S7**

**S6 Derivations** **S8**

S6.1 Proofs for Appendix E.2.2 . . . . . . . . . . . . . . . . . . . . . . . . . . S8

### S1 Application of Algorithm 1 to Simulated Data

The 3D simulations to test Algorithm 1 are described in section 2.3.1 of the main text. The results (shown in Figure S1) are similar to those of the simulations in the main text.

In order to evaluate how the methods compare as the variance changes we generate 1000 realizations (for each realization we generate 50 subjects) and change the variance (which is constant over the image) such that $\frac{1}{\sigma}$ takes values in $\{0.2, 0.4, \dots, 1.4\}$. The results are plotted in right column of Figure S1.

### S2 GLM simulations

The 3D simulations to test Algorithm 3 are described in section 2.3.3 of the main text. In order to approximately match the power of the one-sample simulations, in model (4) we take $\mu$ to have a peak value of 0.5822. The power is only ever approximately the

*Department of Statistics, University of Oxford, Oxford, OX1 3LB, UK

†Oxford Big Data Institute, Li Ka Shing Centre for Health Information and Discovery, Nuffield Department of Population Health, University of Oxford, Oxford, OX3 7LF, UK

‡Wellcome Centre for Integrative Neuroimaging, FMRIB, Nuffield Department of Clinical Neurosciences, University of Oxford, Oxford, OX3 9DU, UK

§Department of Statistics, University of Warwick, Coventry, CV4 7AL, UK

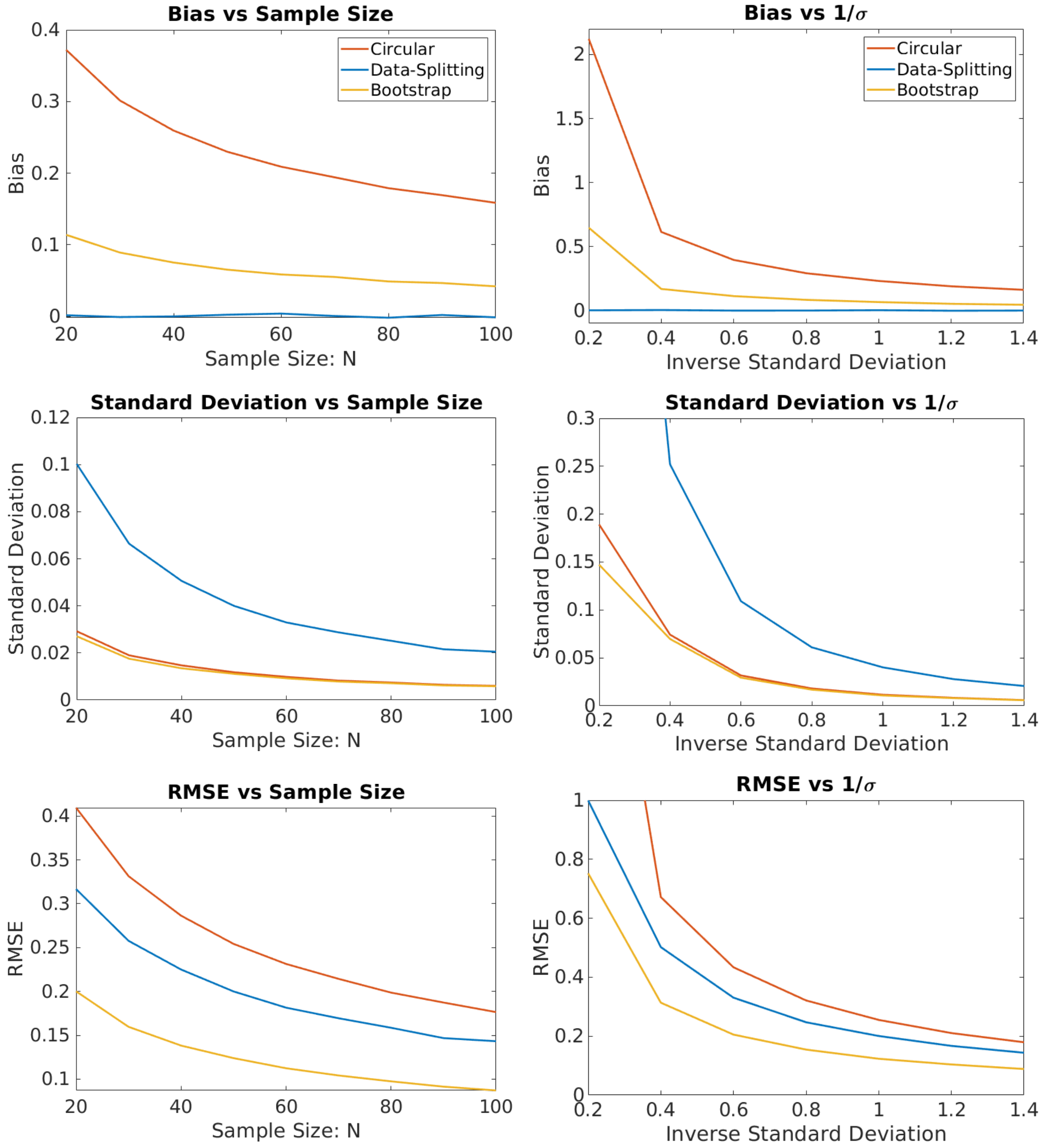


Figure S1: Evaluation as sample size and variance change of bias correction for peaks of the mean (Algorithm 1) on simulated data generated as described in Section 2.3.1. Left column looks at how the measures change with sample size (where the underlying signal and variance are fixed). Right column takes the number of subjects to be 50 and looks at how the measures change with the variance. Each plot shows the bias (top), standard deviation (middle) and RMSE (bottom) calculated over 1,000 realisations. By the overall measure of RMSE, the bootstrap method performs the best.

same as it changes (for the one-sample $t$-statistic versus the $F$-statistic) over sample size and thresholding levels (thus it depends on the FWHM of the noise process). In order to derive the power for model (4) we need

$$f_p^2 = \left(\frac{c^T\beta}{c^T(\mathbb{E}[X^TX])^{-1}c}\right)^2 = \beta^2$$

where $X$ is the model design matrix. The second equality holds as $\mathbb{E}(X^TX)$ is the identity matrix here as the random variables $x$ are mean 0 variance 1 and are independent of the intercept term (as that's just a constant). This yields a corresponding population $R^2 = \frac{\beta^2}{1+\beta^2}$ and allows us to determine the $\beta$ value required to attain a certain level of power, see Appendix E.

The results (shown in Figures S2 and S3) are similar to those of the simulations in the main text. In Figure S2 the bootstrap requires a slightly larger number of subjects (relative to the one sample simulations) before the RMSE drops below that of data-splitting. This is likely because the $R^2$ is a rather complicated function of the subject images and so the bootstrap needs a larger sample size in order to be as effective.

### S3 Additional Simulations for Estimating the Mean at Cohen's $d$ peaks

Figure S4 plots graphs in the same setting as the graphs in Figure 7 (right column) but take $N = 100$ instead of $N = 50$ in order to illustrate that the bootstrap improves (relative to data-splitting) in terms of RMSE for a larger number of subjects.

### S4 Application of Algorithm 1 to fMRI Data

We implemented Algorithm 1, using a threshold of 1.2 % BOLD on the UK biobank fMRI data to obtain barplots, boxplots and graphs which can be interpreted in the same manner as the ones in the main text. See Figures S5 and S6.

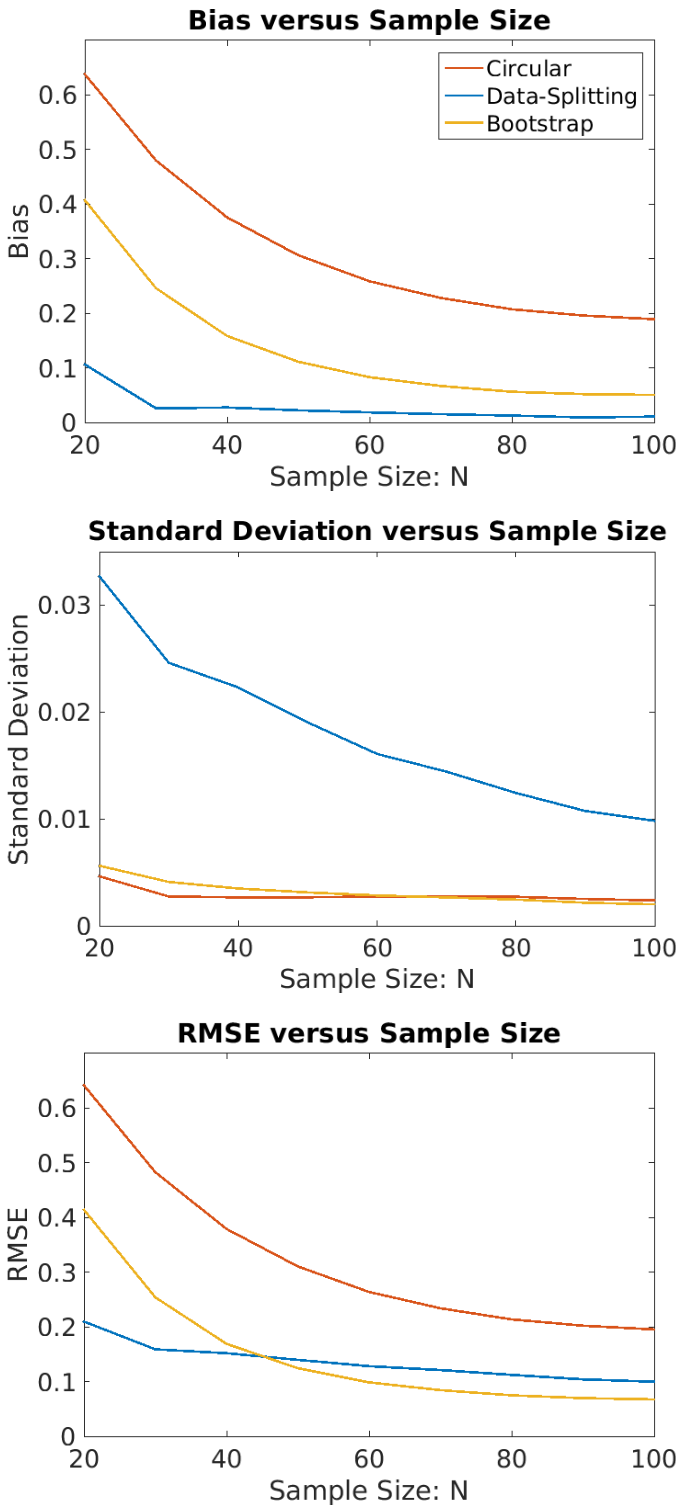


Figure S2: Evaluation of bias correction for $R^2$ peaks (Algorithm 3) as the number of subjects increases on simulated data generated as described in Section S2 (for a peak effect size of $\mu = 0.5822$). Each plot shows the bias (top), standard deviation (middle) and RMSE (bottom) averaged over 1000 realisations, for samples of size $N = 20, 30, \ldots, 100$. By the overall measure of RMSE, the bootstrap method performs the best so long as the sample size is sufficiently large.

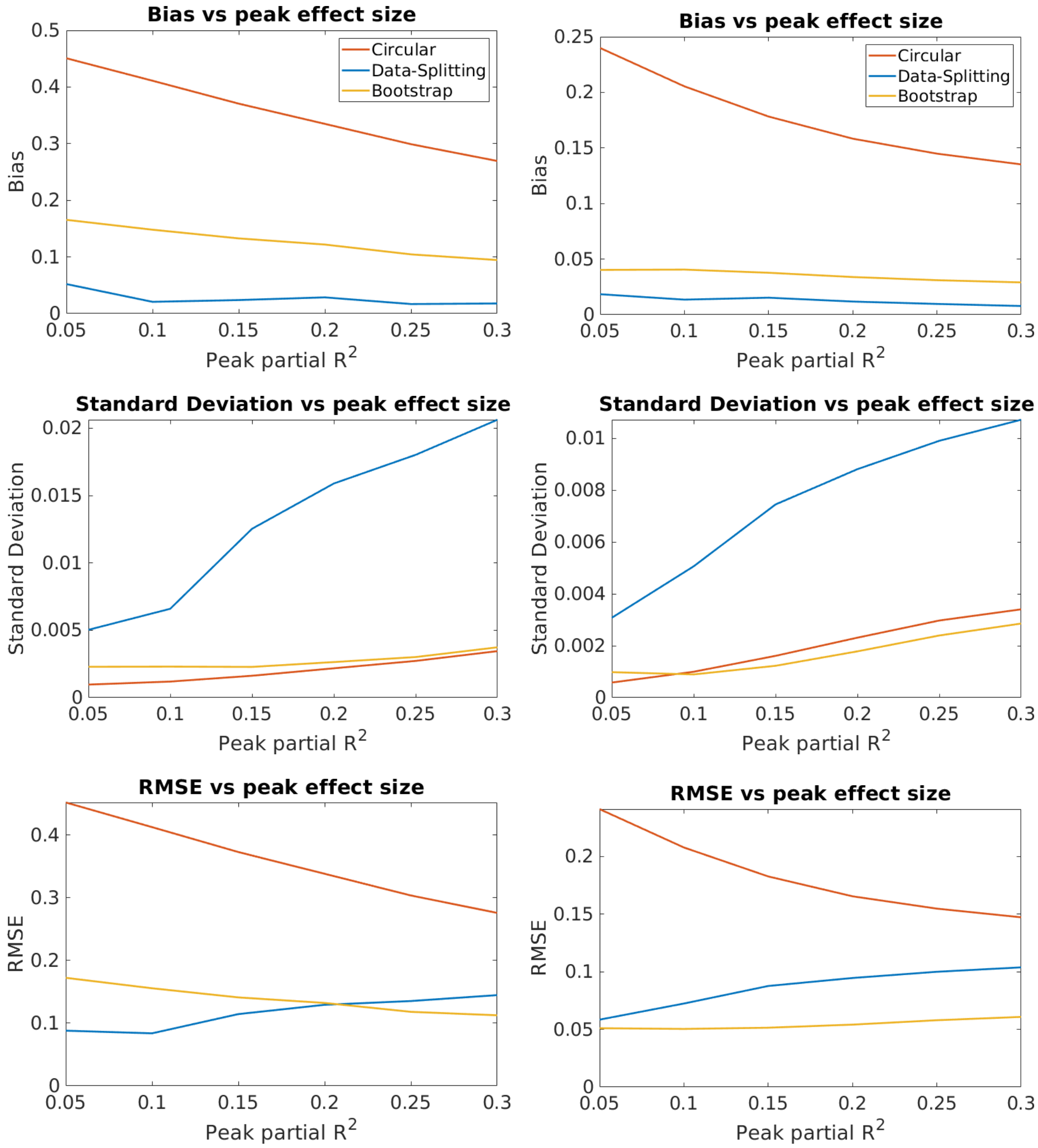


Figure S3: Evaluation as the variance (measured via the peak effect size) changes of bias correction for peaks of $R^2$ (Algorithm 1) on simulated data generated as described in Section 2.3.1. Left column takes $N = 50$ and the right column takes $N = 100$. Each plot shows the bias (top), standard deviation (middle) and RMSE (bottom) calculated over 1,000 realisations. Smaller effect sizes require a larger number of subjects before the bootstrap outperforms data-splitting in terms of RMSE.

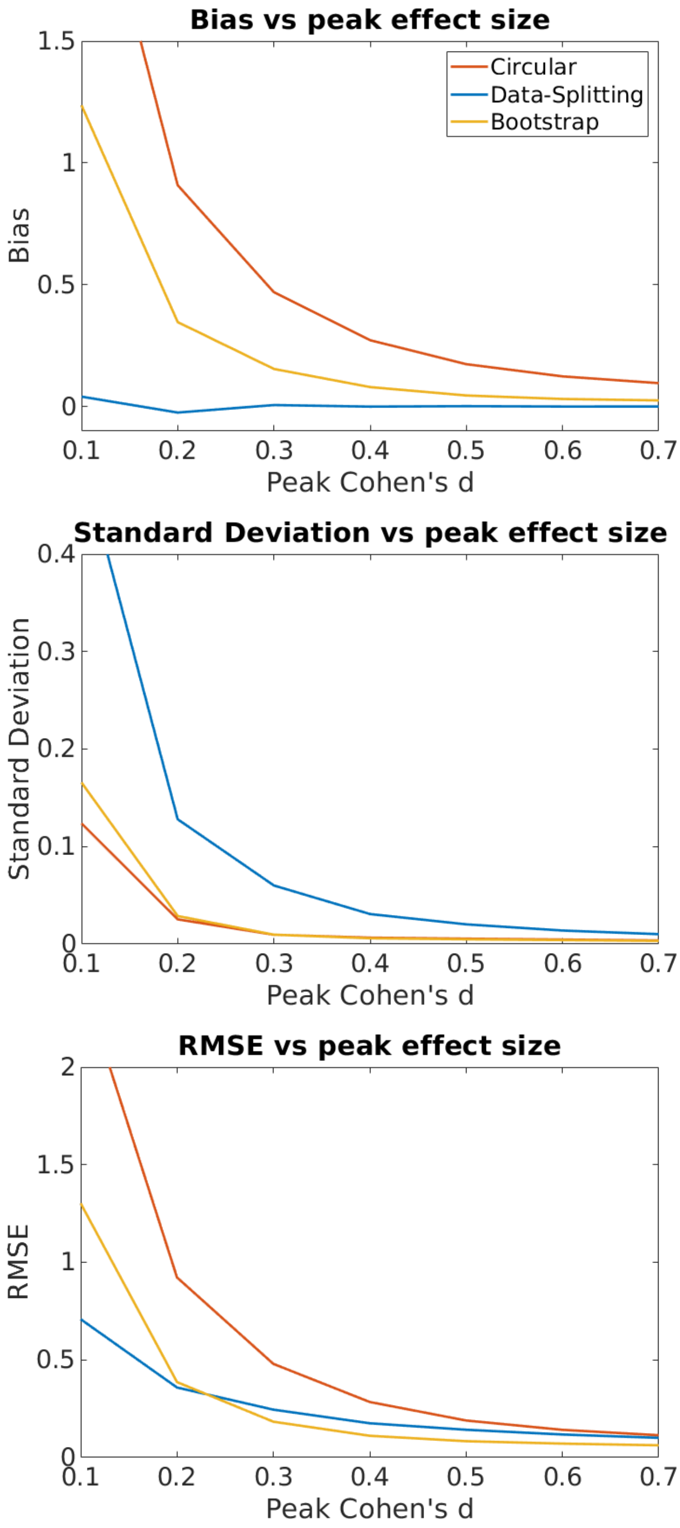


Figure S4: Evaluation as the variance (measured via the peak effect size) changes of bias correction for the %BOLD mean at locations of Cohen's $d$ peaks (Algorithm 2) on simulated data. These graphs are in the same setting as the graphs in Figure 7 (right column) but take $N = 100$ instead of $N = 50$ in order to illustrate that the bootstrap improves (relative to data-splitting) in terms of RMSE for a larger number of subjects. This supports the trend shown in Figure 7 (left column).

### S5 Comparing the Bootstrap and Circular Inference at top peaks

The peak locations found by the bootstrap approach and circular inference are the same. As such we can directly compare the bias/RMSE at the location of the $n$th largest maxima. We have done this in the graph below for the top 15 maxima for the fMRI and VBM datasets. In order to compute these graphs for each $n$ we found the $n$th largest peak in the effect size image ($t$-statistic or partial $R^2$). For instance taking $n = 1$ gives us the maximum, $n = 2$ the second largest peak etc. Given a number of subjects $N$ and a peak rank $n$, we get $G_N = \lfloor 4940/N \rfloor$ peaks of rank $n$ and obtain estimates $\hat{\theta}_1^n, \dots \hat{\theta}_{G_N}^n$ for the values of the underlying effects $(\theta_1^n, \dots \theta_{G_N}^n)$ at the locations of these peaks (using circular inference and the bootstrap methods). As in the main text (Section 2.5) the underlying effects take different values since the locations are different. As such for $k = 1, \dots, G_N$ we compare the differences $\hat{\theta}_k^n - \theta_k^n$ to 0 (where the $\theta_k^n$ are computed using the 4000 subject held out ground truth) and compute

$$\text{Bias}_n = \frac{1}{G_N}\sum_{k=1}^{G_N}\left(\hat{\theta}_k^n - \theta_k^n\right) \quad \text{and} \quad \text{RMSE}_n = \left(\frac{1}{G_N}\sum_{k=1}^{G_N}(\hat{\theta}_k^n - \theta_k^n)^2\right)^{1/2}.$$

We have plotted these quantities against $n$ in Figures S7 and S8. We note that the graphs are somewhat variable because we have used the 4940 subjects to calculate them meaning that for each $N = 50, 100, 150$ we only have $98, 49, 32$ (respectively) peaks for each $n$. From the graphs we see the bootstrap significantly outperforms circular inference (at all peak ranks). The bootstrap estimates are relatively unbiased and have low RMSE whereas the circular estimates are substantially positively biased and have a larger RMSE.

### S6 Derivations

#### S6.1 Proofs for Appendix E.2.2

Under the framework of Appendix E.2.2, we have:

**Proposition S6.1.**

$$\frac{1}{N} X_N^T X_N \xrightarrow{a.s.} \mathbb{E}\big[x_1 x_1^T\big],$$

$$\hat{\beta}_N \xrightarrow{a.s.} \beta$$

*and*

$$\hat{\sigma}_N^2 \xrightarrow{a.s.} \sigma$$

*as* $N \longrightarrow \infty$*, where* $\xrightarrow{a.s.}$ *denotes pointwise almost sure convergence.*

*Proof.*

$$\frac{1}{N} X_N^T X_N = \frac{1}{N} \sum_{k=1}^{N} x_k x_k^T \xrightarrow{a.s.} \mathbb{E}\big[x_1 x_1^T\big]$$

as $N \longrightarrow \infty$ by the strong law of large numbers (SLLN) as the variance of the $x_i$ is finite. As such

$$\hat{\beta}_N = (X_N^T X_N)^{-1} X_N^T Y_N = \beta + \left(\frac{1}{N} X_N^T X_N\right)^{-1} \frac{1}{N} X_N^T \epsilon_N \xrightarrow{a.s.} \beta$$

by applying Slutsky since the SLLN implies that $X_N^T \epsilon_N / N = \sum_{k=1}^N \epsilon_k x_k / N \xrightarrow{a.s.} 0$ as $N \longrightarrow \infty$ since by independence the expectation is 0. Note Cauchy-Schwartz and the finite variance conditions are used here in order to show that the expected absolute first moment is finite and thereby justify the convergence.

It follows that for every $\eta > 0$ there is some large enough $N$ such that $\|\hat{\beta}_N - \beta\|^2 < \eta$ and in particular for large enough $N$,

$$\frac{1}{N-p} \left\| X_N \beta - X_N \hat{\beta}_N \right\|^2 = \frac{1}{N-p} \sum_{i=1}^{N} \left( x_i^T \hat{\beta}_N - x_i^T \beta \right)^2 \leq \frac{\eta}{N-p} \sum_{i=1}^{N} \|x_i^T\|^2 \longrightarrow \eta \mathbb{E} \|x_i^T\|^2$$

which tends to zero as $N \longrightarrow \infty$ since $\eta$ can be made arbitrarily small. Also, for any $\eta$ and large enough $N$, by Cauchy-Schwartz,

$$\frac{1}{N-p} \left| (Y_N - X_n \beta)^T (X_N \beta - X_N \hat{\beta}) \right| = \frac{1}{N-p} \sum_i \epsilon_i | x_i^T \beta - x_i^T \hat{\beta}_N | \leq \frac{\eta^{1/2}}{N-p} \sum_i \epsilon_i \|x_i^T\|$$

so this converges to zero as $N \longrightarrow \infty$ by the SLLN.

$$\begin{aligned}
\hat{\sigma}_N^2 &= \frac{1}{N-p} \|Y_N - X_N \hat{\beta}_N\|^2 \\
&= \frac{1}{N-p} \|Y_N - X_N \beta\|^2 + \frac{1}{N-p} \|X_N \beta - X_N \hat{\beta}\|^2 + \frac{2}{N-p} (Y_N - X_N \beta)^T (X_N \beta - X_N \hat{\beta}_N).
\end{aligned}$$

So $\hat{\sigma}_N^2 \xrightarrow{a.s.} \sigma^2$ as $N \longrightarrow \infty$ since the last two terms tend to 0 and the first equals $\frac{1}{N-p} \sum_i^N \epsilon_i^2$ and converges by the SLLN. □

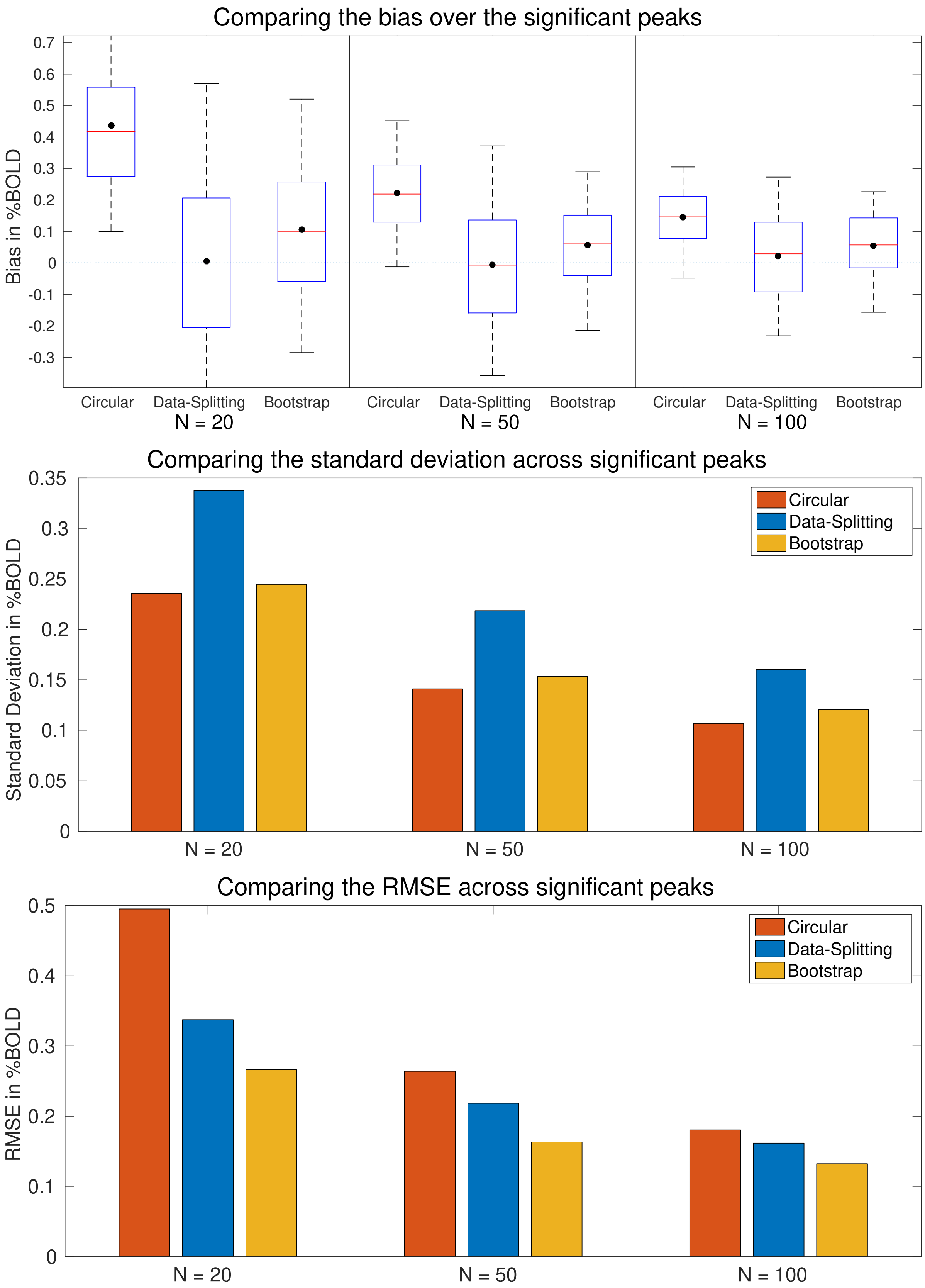


Figure S5: Comparison of estimates for the sample mean via Algorithm 1. Bias (top), variance (middle), and MSE (bottom) are shown for $N = 20, 50$ and 100 sample sizes, based on $G_N$ samples. The bootstrap estimates display some bias but have lower MSE than the data-splitting estimates.

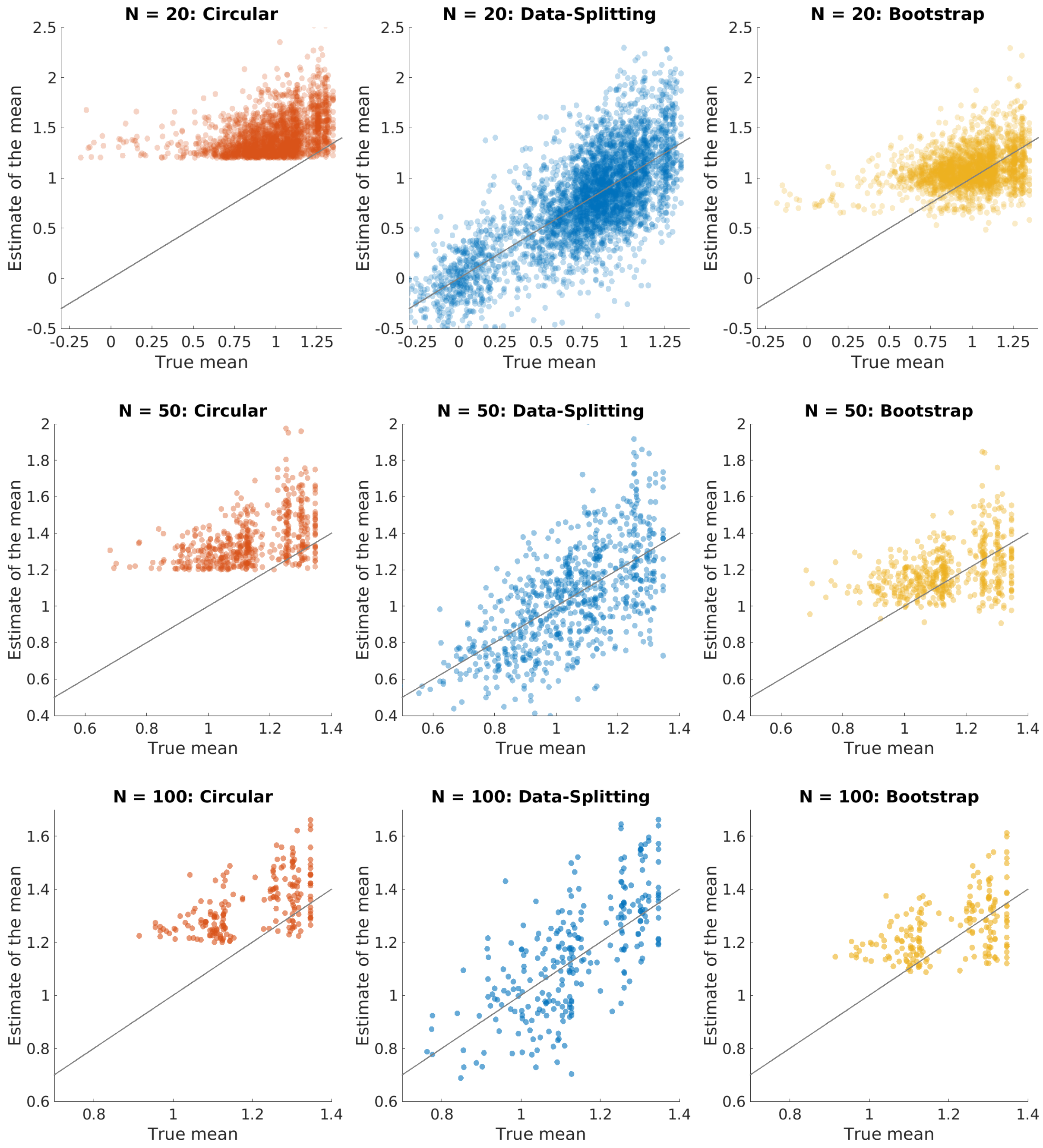


Figure S6: Plots of estimated versus true value of the sample mean, for circular (left), data-splitting (middle), and bootstrap (right). Plots show all peaks found over the $G_N$ samples for each sample size, $N = 20, 50, 100$ (top to bottom). For each peak the true sample mean is obtained at that location from the held-out 4000 subject mean image. Note that the number of peaks and their locations are the same for circular inference and the bootstrap but are different for data-splitting as it uses the first half of the subjects in order to determine significant peaks. From these plots we see that the data-splitting estimates are unbiased but are more variable than the bootstrap and circular estimates.

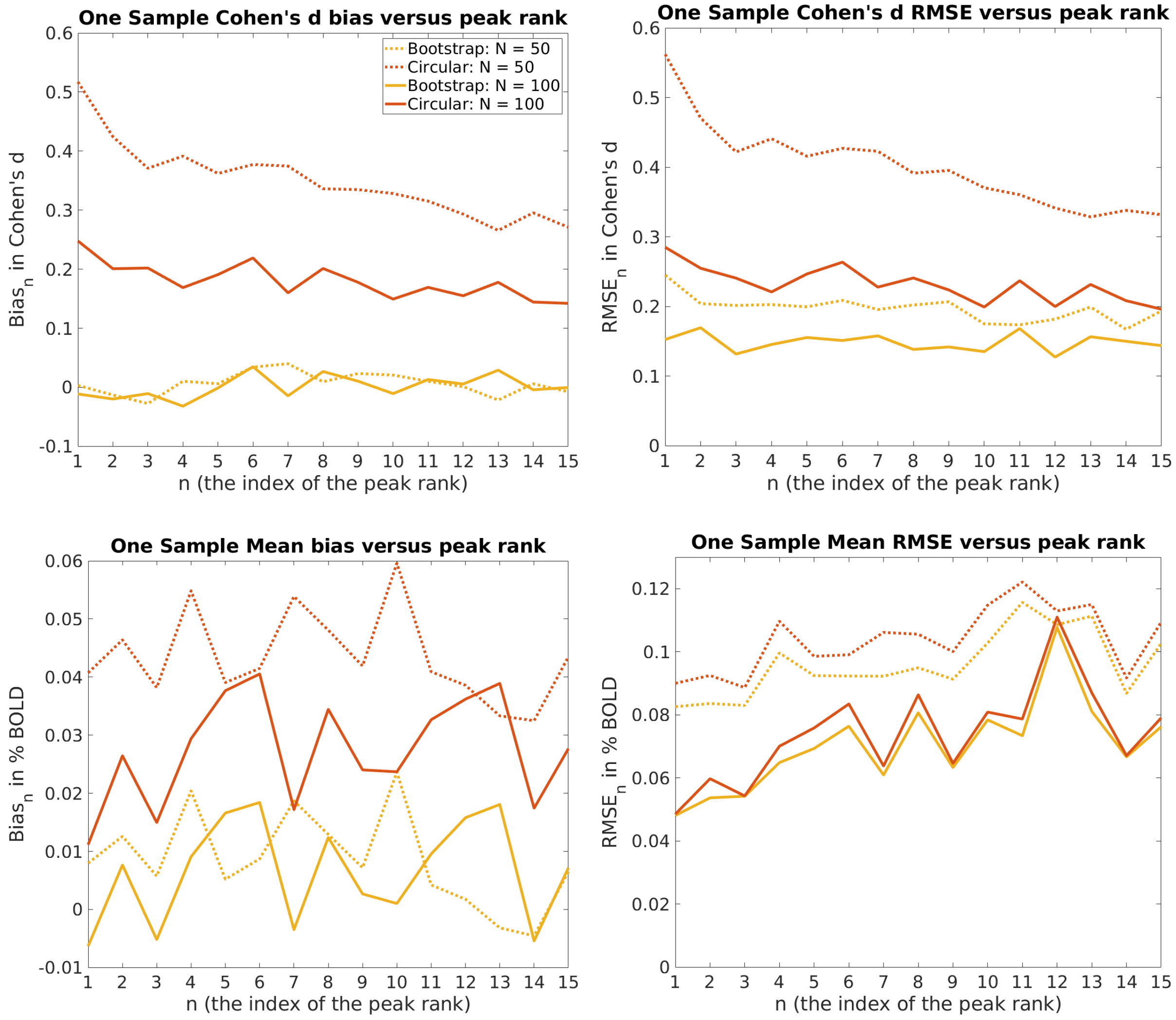


Figure S7: Comparison of estimates, at the top peaks of the t-statistic of the one-sample mean and Cohen's $d$, for task fMRI images. Here the bootstrap estimates are calculated using Algorithm 2. Plots of the bias are shown on the left and plots of the RMSE are shown on the right. The bootstrap curves (shown in yellow) always lie below those of the circular curves (shown in red). In particular they show that at each peak rank the bootstrap has low bias and low RMSE relative to circular inference.

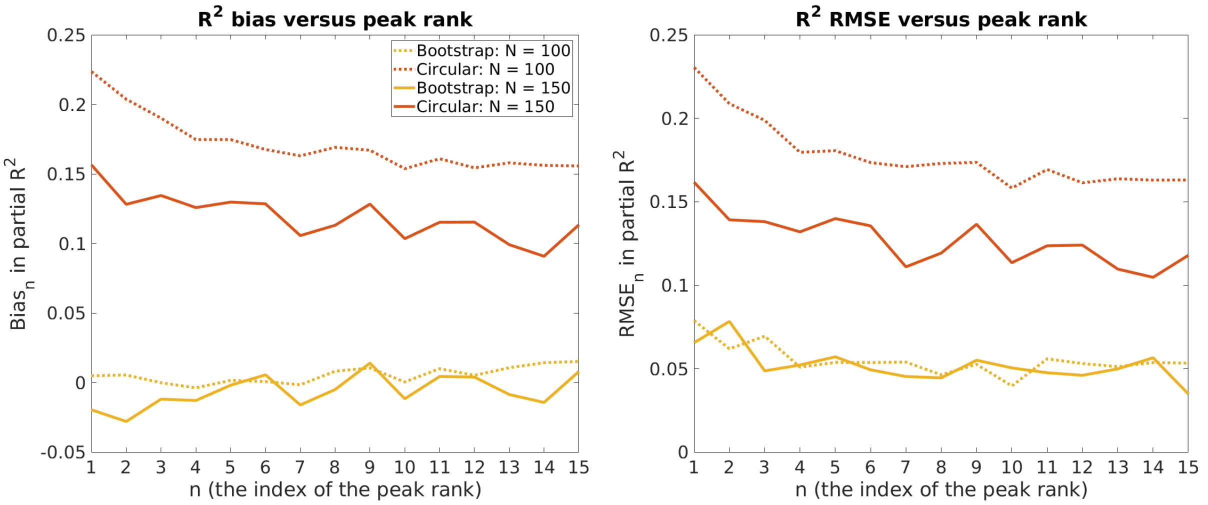


Figure S8: Comparison of estimates at the top peak ranks for the partial $R^2$ for age obtained using a GLM regression on VBM data. Plots of the bias are shown on the left and plots of the RMSE are shown on the right. The bootstrap curves (shown in yellow) always lie below those of the circular curves (shown in red). In particular they show that at each peak rank the bootstrap has low bias and low RMSE relative to circular inference.
